# Airway secretory cell-derived p63^+^ progenitors contribute to alveolar regeneration after sterile lung injury

**DOI:** 10.1101/2023.02.27.530122

**Authors:** Zan Lv, Zixin Liu, Kuo Liu, Wenjuan Pu, Yan Li, Huan Zhao, Ying Xi, Andrew E. Vaughan, Astrid Gillich, Bin Zhou

## Abstract

Lung injury activates epithelial stem or progenitor cells for alveolar repair and regeneration. However, the origin and fate of injury-induced progenitors are poorly defined. Here, we report that p63-expressing progenitors emerge upon bleomycin-induced lung injury. These p63^+^ progenitors proliferate rapidly and differentiate into alveolar type 1 (AT1) and type 2 (AT2) cells through distinct trajectories. Dual recombinase-mediated sequential genetic lineage tracing reveals that p63^+^ progenitors originate from airway secretory cells and subsequently generate alveolar cells. Functionally, p63 activation is required for efficient alveolar regeneration from secretory cells. Our study identifies a secretory cell-derived p63^+^ progenitor that contributes to alveolar repair, indicating a potential therapeutic avenue for lung regeneration after injury.

## Introduction

Efficient gas exchange in the lung depends on the delicate organization, integrity and function of diverse types of epithelial cells. Lung epithelial cells are located in two structurally distinct compartments: the conducting airways and the alveoli, the sites of gas exchange. Following lung injury caused by inhaled toxins, respiratory viruses or bacteria, stem cells or progenitors are activated and generate new epithelial cells for compartment-specific repair^1^. For example, in the airway, local epithelial progenitors contribute to regeneration after injury^2–5^. In the setting of alveolar injury, surfactant-secreting alveolar type 2 (AT2) epithelial cells contribute to alveolar renewal and repair by proliferating and differentiating into thin alveolar type 1 (AT1) cells that mediate gas exchange^6,7^. More specifically, a subpopulation of AT2 cells expressing *Axin2*, a transcriptional target of Wnt signaling, exhibits stem cell activity^8^. These Wnt-responsive AT2 cells proliferate and differentiate into AT1 cells in the distal lung upon injury^9^. In addition, a unique stem cell population residing at the bronchioalveolar junction acts as progenitors for both distal airway and alveolar epithelial cells after naphthalene and bleomycin-induced lung injury, respectively^10–12^. During severe injury induced by influenza, lineage-negative epithelial stem/progenitor cells (LNEP) present in the distal lung mobilize to regenerate alveolar epithelial cells^13,14^. A distinct subpopulation of club-like LNEPs marked by high H2-K1 expression also plays a role in alveolar repair. Upon transplantation, H2-K1^high^ cells differentiate into alveolar cells in injured lungs^15^. However, direct *in vivo* genetic lineage tracing evidence to demonstrate the endogenous fate and function of H2-K1^high^ progenitors is lacking^15^. Conversely, p63^+^ LNEPs represent a rare population of intrapulmonary basal-like cells that expand dramatically upon influenza injury but rarely give rise to alveolar cell types, instead representing a dysplastic repair response^16–18^. In spite of these advances, the origins and lineage relationships of these various progenitor populations remain unclear, and whether their relative fate potential is dictated by the nature of the injury is similarly understudied.

Regeneration of alveoli from different types of stem/progenitor cells involves their stepwise differentiation to mature alveolar epithelial cells. However, the cellular and molecular programs for these cell fate transitions have yet to be fully elucidated. A recent study reports that inhibition of Notch signaling triggers the conversion of secretory cells into AT2 cell lineages after lung injury, and this conversion involves two sequential stages including a *Flst1*^+^ secretory cell stage^19^. Additionally, distal airway epithelial cells are heterogeneous and may contain diverse stem cell states including the newly identified *SFTPC*^+^ *SCGB3A2*^+^ alveolar type-0 (AT0) and respiratory airway secretory (RAS) cells that may contribute to alveolar regeneration in humans^20,21^. After influenza virus infection or bleomycin treatment, p63^+^ progenitor cells are induced in damaged lung parenchyma^13,22^. These p63^+^ cells upregulate additional basal cell markers (Krt5), migrate to injured alveoli, and form bronchiole-like lumen structures, but do not appear to contribute appreciably to alveolar regeneration^22^. Indeed, most studies indicate that pre-existing p63-labeled basal cells rarely generate alveolar epithelial cells after H1N1 influenza infection, though some groups, by inducing *p63-CreERT2* lineage labeling during injury, have suggested that the cells bear somewhat higher potential to contribute to alveolar repair^23^. Intriguingly, initial studies showing that p63^+^ cells rarely generate alveolar epithelium after influenza also demonstrated that these cells had greater potential to generate AT2 cells after injury with bleomycin – up to ~1/3 of Krt5 lineage-labeled cells ultimately express AT2 markers^13^. In spite of these advances, the origins, fates, and functions of p63^+^ cells during lung repair remain enigmatic. The relationship between secretory cells and injury-induced progenitors, as well as why the fate potential of these cells seems different in various injury models, needs to be elucidated.

In this study, we identify p63-expressing progenitors that emerge in alveoli following bleomycin lung injury. Fate mapping of these p63^+^ cells reveals a contribution to AT1 and AT2 cells after injury resolution. Single-cell RNA-sequencing (scRNA-seq) suggests that the p63^+^ progenitors contribute to AT1 and AT2 cells through distinct differentiation routes. Furthermore, genetic lineage tracing using dual recombinases demonstrates that the injury-induced p63^+^ progenitors are derived from airway secretory cells. Notably, p63 is not only a marker for progenitors but also functionally required for the activation, proliferation, and reprogramming of secretory cells to AT2 cells. These findings link distal airway progenitors to alveolar epithelial repair and regeneration through an injury-induced progenitor.

## Results

### Injury-induced p63^+^ Cells Proliferate and Differentiate into Alveolar Cells

To genetically label p63-expressing cells *in vivo*, we generated a *p63-CreER* knock-in mouse line, in which *2A-CreER* sequence was inserted into the *p63* gene locus (Extended Data Fig. 1a). By crossing the line with the Cre-responsive reporter *Rosa26-tdTomato* (*R26-tdT*)^24^, we found that *p63-CreER* efficiently labels p63^+^ cells after tamoxifen (Tam) treatment (Extended Data Fig. 1b,c). We detected lineage labeling of basal cells in the trachea and rare cells in the lung (presumably p63^+^ LNEPs) as expected^13,14^. There was virtually no leakiness of Cre activity without Tam treatment (Extended Data Fig. 1c). To determine if p63^+^ cells expand in lungs upon injury, we administered Tam to *p63-CreER;R26-tdT* mice and induced alveolar injury two weeks later by intratracheal instillation of bleomycin (Extended Data Fig. 1d) as previously described^11^. We observed efficient lineage labeling of tracheal basal cells but only rare tdT^+^ cells in the lung (Extended Data Fig. 1e-f). This suggests that pre-existing intrapulmonary p63^+^ LNEPs that expand upon influenza injury do not expand upon sterile injury with bleomycin^16,18,25^. By using *K5-2A-CreER;R26-tdT* mice, we did not detect significant contribution of pre-labeled K5^+^ cells to alveoli after injury (Extended Data Fig. 1g-l).

To label injury-activated p63^+^ cells, we treated mice with Tam after bleomycin administration (Figure 1a). Whole-mount fluorescence images of lungs collected at 14 days after bleomycin treatment (Bleo D14) revealed abundant tdT^+^ signals in injured lungs (Figure 1b). Light-sheet imaging of tdT signals in cleared left lobes of injured lungs showed that most tdT^+^ cells were arranged in patches (Figure 1c, e). Immunostaining for tdT with the airway epithelial cell marker CC10 and the alveolar epithelial cell marker AGER (RAGE) on lung sections showed that the tdT^+^ patches were located in alveolar regions close to bronchioles (Figure 1d, f). At Bleo D14, only a subset of tdT^+^ cells expressed p63 (Figure 1g), suggesting that p63 is relatively quickly downregulated as injury resolves. Among tdT^+^ cells, we also noticed some p63 low-expressing cells that began to exhibit weak expression of AT2 cell marker, Surfactant Protein-C (SPC, Figure 1h), indicating transitional cell status of simultaneous p63 and SPC expression. Immunostaining for additional lineage markers revealed that more than half (51%) of tdT^+^ cells express alveolar epithelial cell markers (SPC or AGER) (Figure 1i), indicating that a substantial portion of p63-derived cells adopt alveolar epithelial cell fate during injury repair.

**Figure 1.**
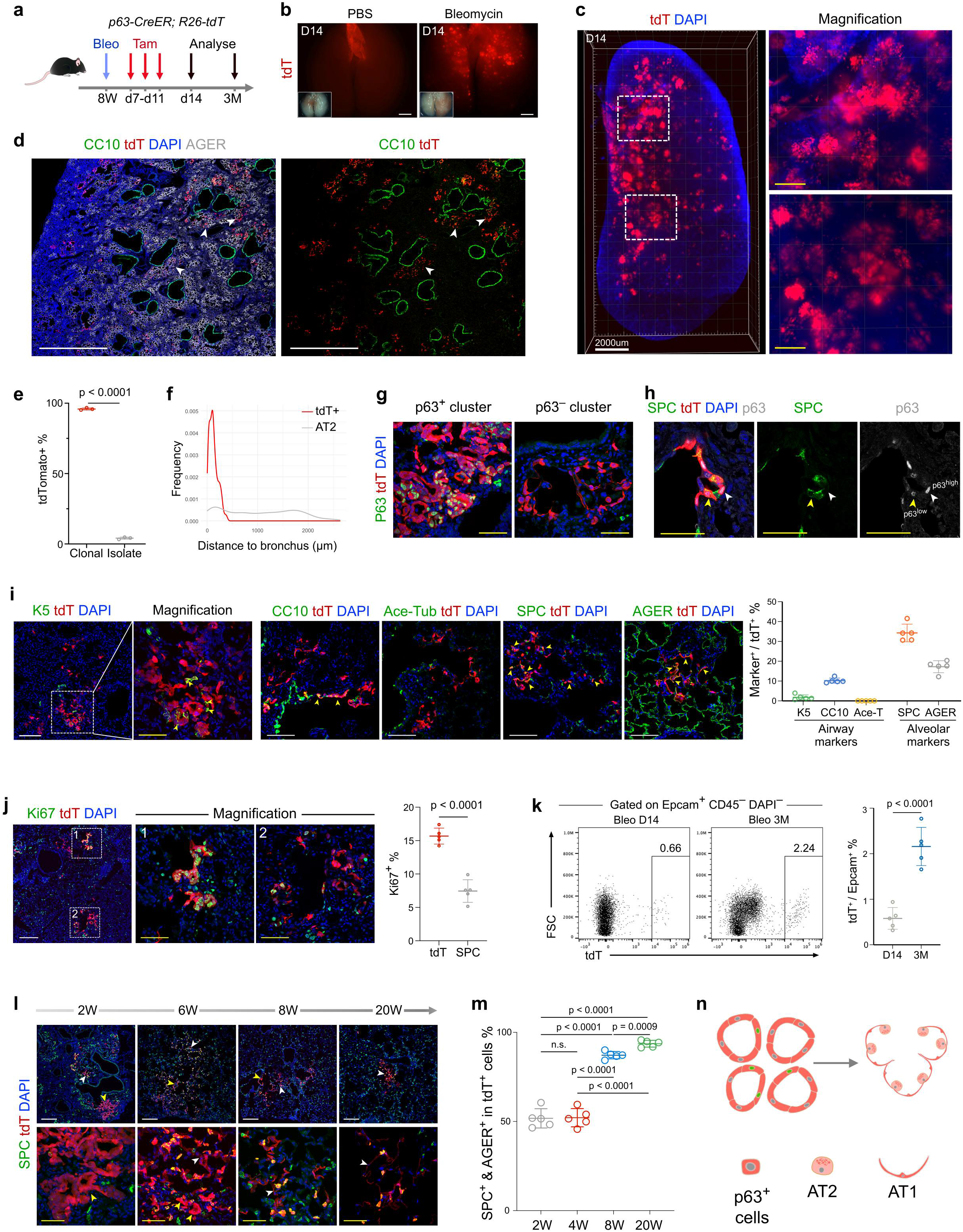
Injury-induced p63^+^ progenitors contribute to alveolar regeneration. **a**, A schematic showing the experimental design. **b**, Whole-mount fluorescent views of PBS or bleomycin-treated lungs. Inserts indicate bright-field images. Scale bars, 2mm. **c**, Reconstructed 3D light-sheet microscopic image of the injured lungs on day 14 after bleomycin treatment. Scale bar, white, 2mm; yellow, 500 μm. **d**, Immunostaining for CC10, tdT, and AGER on injured lungs of *p63-CreER;R26-tdT* mice at 14 dpi. Arrowheads indicate tdT^+^ cells residing in proximity to CC10^+^ bronchioles. Scale bar, 1mm. **e**, Quantification of the percentage of tdT^+^ cells that are clonal or isolated. **f**, Quantification of the distance of tdT^+^ cells and SPC^+^ cells to bronchioles. **g**, Immunostaining for p63 and tdT on lung sections. Scale bar, 50 μm. **h**, Immunostaining for SPC, tdT, and p63 on lung sections. Yellow arrowhead, p63^low^SPC^+^ cells; white arrowhead, p63^high^SPC^−^ cells. Scale bars, 50 μm. **i**, Immunostaining for different lineage markers and tdT on lung sections. Yellow arrowheads, marker^+^tdT^+^ cells; Right panel shows quantification of the percentage of tdT^+^ cells expressing K5, CC10, acetylated-tubulin, SPC, or AGER in the injured lungs. n = 5 mice for each group. Scale bar, white, 100 μm; yellow, 50 μm. **j**, Immunostaining for Ki67 and tdT on lung sections. Right panel shows quantification of the percentage of tdT^+^ or SPC^+^ cells expressing Ki67. Scale bar, white, 200 μm; yellow, 50 μm. **k**, Flow cytometric analysis of the percentage of Epcam^+^ cells expressing tdT. n = 5 mice for each group. **l**, Immunostaining for SPC and tdT on lung sections collected at indicated time points. Yellow arrowheads, honey-comb structure; white arrowheads, alveolar structure. Scale bar, white, 200 μm; yellow, 50 μm. **m**, Quantification of the percentage of tdT^+^ cells expressing SPC or AGER at indicated time points. n = 5 mice for each group. **n**, A cartoon image showing contributeion of p63^+^ cells to alveolar epithelial cells after lung injury.

We examined proliferation of tdT^+^ cells using the cell cycle marker Ki67 and found that a subset of tdT^+^ cells express Ki67 (Figure 1j). The percentage of cells expressing Ki67 was significantly higher in tdT^+^ cells than tdT^−^ SPC^+^ AT2 cells (Figure 1j). Long-term tracing of p63-activated cells showed a significant increase of the percentage of epithelial cells expressing tdT in lungs collected at 3 months after bleomycin treatment (Bleo 3M) compared with Bleo D14 (Figure 1k), suggesting a robust expansion of p63-labeled cells during repair. We next examined the differentiation process of p63^+^ progenitors by collecting samples at 2 weeks (W), 6W, 8W, and 20W after bleomycin treatment (Figure 1l). We observed that the shape of tdT^+^ patches gradually changed from honeycomb-like to alveolar sac-like and a subset of tdT^+^ cells acquired a flattened morphology over time (Figure 1l). The percentage of lineage-labeled cells that express alveolar markers increased from 2W to 20W (Figure 1m), demonstrating that the majority of injury-activated p63 cells adopt alveolar epithelial cell fates. Taken together, we identified injury-induced p63^+^ cells that self-renew and differentiate into alveolar epithelial cells after injury (Figure 1n), supporting p63^+^ cells as progenitors for alveolar repair and regeneration.

### Transitional Cell States of p63^+^ Progenitors in Alveolar Differentiation

To understand how p63^+^ progenitors progress to generate alveolar epithelial cells, we performed single-cell RNA sequencing (scRNA-seq) of p63-traced cells isolated from *p63-CreER;R26-tdT* mouse lungs at 14 days (D14), 2 months (2M), and 5 months (5M) after bleomycin treatment (Figure 2a, Extended Data Fig. 2a, b). Uniform manifold approximation and projection (UMAP) identified two major cell clusters including tdT^+^ epithelial cells and tdT^low^ F4/80^+^ macrophages (Extended Data Fig. 2c). We eliminated airway epithelial cells and immune cells and re-clustered the remaining cells (Figure 2b). The p63-derived cells consisted of four cell clusters expressing p63, p63^high^ cell 1-4 (p63-1, p63-2, p63-3, p63-4), AT1, AT2, and five intermediate clusters (Krt8^+^, Transient 1, Transient 2, Pre-AT2, AT2-to-AT1) of which all expressed tdT with distinct molecular signatures (Figure 2b,c,d). Of note, a subset of p63-derived cells expressed basal cell marker *Krt5*, which was mainly restricted in p63-1 and p63-2 subpopulations (Figure 2c). We localized and characterized the newly identified intermediate cells in injured lung tissues by immunostaining. We detected Aqp3^+^, WFDC1^+^, Scgb3a2^+^, and Krt8^+^ tdT lineage-labeled cells on tissue sections, representing p63^+^ cluster, Transient cell-1, Transient cell-2, Krt8^+^ cells, respectively (Figure 2e), indicating that the cell clusters we identified by scRNA-seq analysis represent *in vivo* cell states.

**Figure 2.**
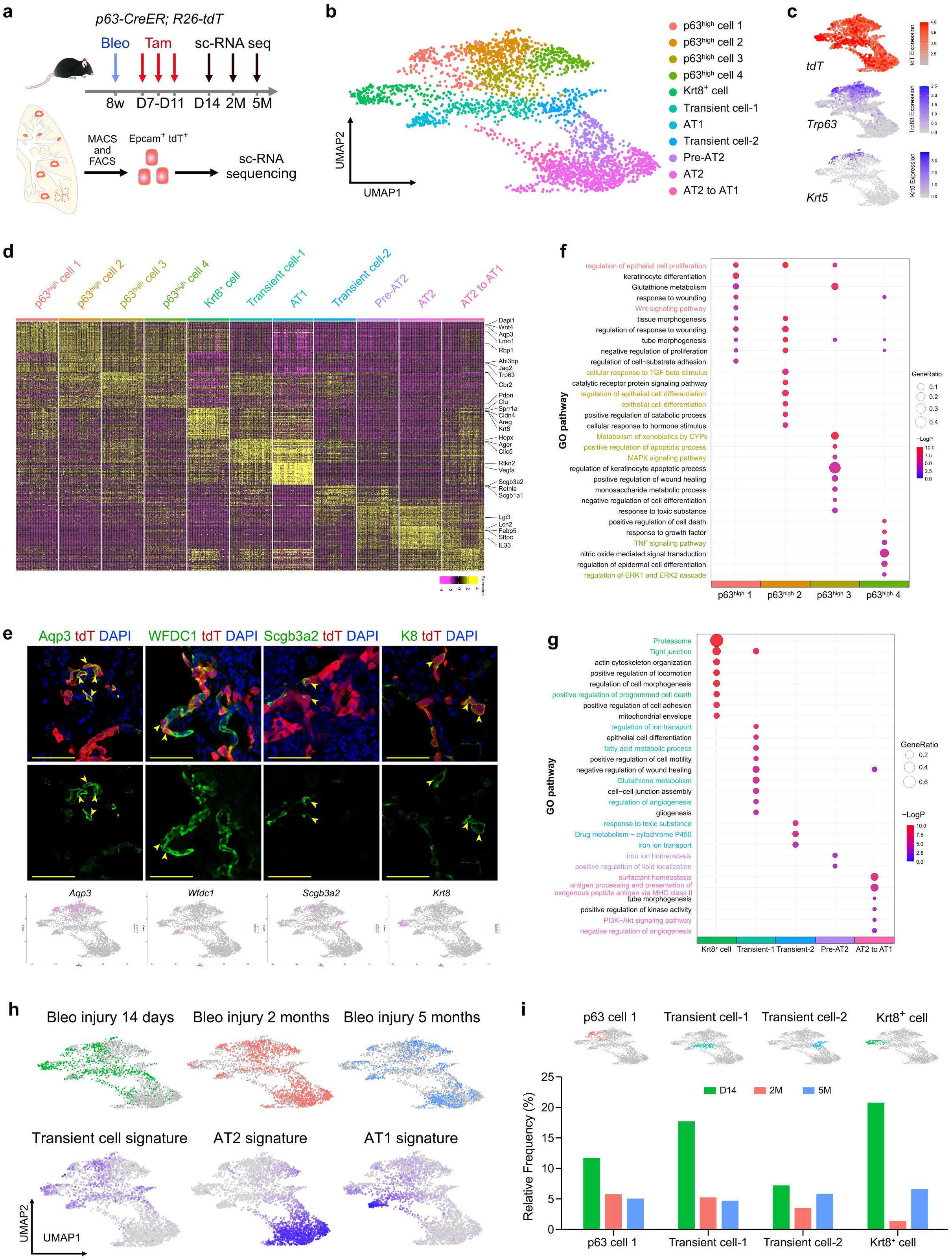
scRNA-seq analysis of p63^+^ progenitors. **a**, A schematic showing the experimental design. Epcam^+^tdT^+^ cells were sorted to perform scRNA-seq at 14 days (D14), 2 months (2M) and 5 months (5M) after bleomycin injury. **b**, UMAP visualization of the epithelial populations in bleomycin-treated murine lungs (n = 2896 cells): P63^high^ cell (n = 1007 cells); Krt8^+^ cell (n = 221 cells); Transient cell-1 (n = 235 cells); AT1 (n = 96 cells); Transient cell-2 (n = 140 cells); Pre-AT2 (n = 241 cells); AT2 (n = 850 cells); AT2-to-AT1 (n = 106 cells). **c**, UMAP plots showing the expression of *tdT* and *p63* in epithelial populations in bleomycin-treated lungs. **d**, Heatmap plot showing the expression of marker genes across different cell subpopulations in bleomycin-treated lungs. **e**, Immunostaining for Aqp3, WFDC1, Scgb3a2, and Krt8 on lung sections collected at 14 dpi. UMAP plots showing the expression of these genes under the staining images. Scale bar, 50 μm. **f**, Enrichment analysis reveals pathways enriched in each p63^high^ cell subpopulation respectively. The web-based tool Metascape was used to determine the adjusted p-value and gene ratio. **g**, Enrichment analysis reveals pathways enriched in Krt8^+^ cell, Transient cell-1, Transient cell-2, Pre-AT2, and AT2-to-AT1 subpopulations, respectively. The web-based tool Metascape was used to determine the adjusted p-value and gene ratio. **h**, UMAP plots showing the sample source for cells (top) and specific gene sets signature (bottom). **i**, UMAP plots showing the selected cell subpopulations (top) and barplot showing the relative frequency of the indicated cell types relative to other cells at the indicated time points after injury (bottom).

Gene ontology analysis showed that the p63-1 cluster was highly enriched in epithelial cell proliferation regulation genes (Ccnd2) and the Wnt signaling pathway (Figure 2f). The other three p63^+^ clusters displayed distinct signatures – response of TGF-β stimulation in the p63-2 cluster, MAPK signaling in the p63-3 cluster, and TNF signaling in the p63-4 cluster (Figure 2f, Extended Data Fig. 2d,f). These clusters were also enriched for differentiation or programmed cell death-associated signatures, indicating that they represent a more differentiated state compared to p63^+^ progenitor cells. As part of the five intermediate clusters, we identified a Krt8^+^ cluster (Extended Data Fig. 2e), which was similar to the previously reported transient state of AT2 to AT1^26,27^. We also found that the Transient 1 cell cluster was enriched for tight junction, cell-cell junction assembly, and glutathione metabolism, whereas the Transient 2 cell cluster was enriched for ‘response to a toxic substance and cytochrome P450’ (Figure 2g). The constitution of these clusters changed as regeneration progressed, as cells with transient cell signatures were enriched at 14 days after injury, whereas mature AT1 and AT2 cells were more abundant at 2 and 5 months post-injury (Figure 2h). This was supported by a decrease in abundance of p63-1, transient 1, transient 2, and K8 cells at 2 and 5 months post-injury (Figure 2i, Extended Data Fig. 2g).

### p63^+^ Progenitors Contribute to AT1 and AT2 Cells through Two Trajectories

Considering the surprising diversity of p63^+^ cells with multiple intermediate states, we next examined the relationships between the clusters and the potential trajectory of p63-derived lineages. RNA velocity analysis showed that the p63-1 cluster contributes to AT1 and AT2 cells through two distinct trajectories (Figure 3a). Specifically, trajectory 1 represents the differentiation of p63-1 cells to AT1 cells through p63-3 cells and Transient 1 cells. Trajectory 2 represents the differentiation of p63-1 cells to AT2 cells through step-wise conversion of p63-3 cells to Transient 2 cells and then Pre-AT2 cells (Figure 3b). We analyzed the transcriptome changes in these two trajectories to identify genes that are transiently activated during the differentiation processes. The genes transiently activated in trajectory 1 were clustered in epithelial cell proliferation, wound healing, and cell-cell junction organization, whereas genes transiently activated in trajectory 2 were clustered in the regulation of angiogenesis, response to hypoxia, and regulation of ATPase activity (Figure 3c,d). By comparing the unique genes that were only activated in trajectory 1 or trajectory 2, we found that genes associated with wound healing and positive regulation of kinase activity were preferentially activated in trajectory 1, whereas genes with roles in angiogenesis and positive regulation of cell adhesion were activated predominantly in trajectory 2 (Figure 3e,f), suggesting that distinct molecular programs regulate these cell fate transitions.

**Figure 3.**
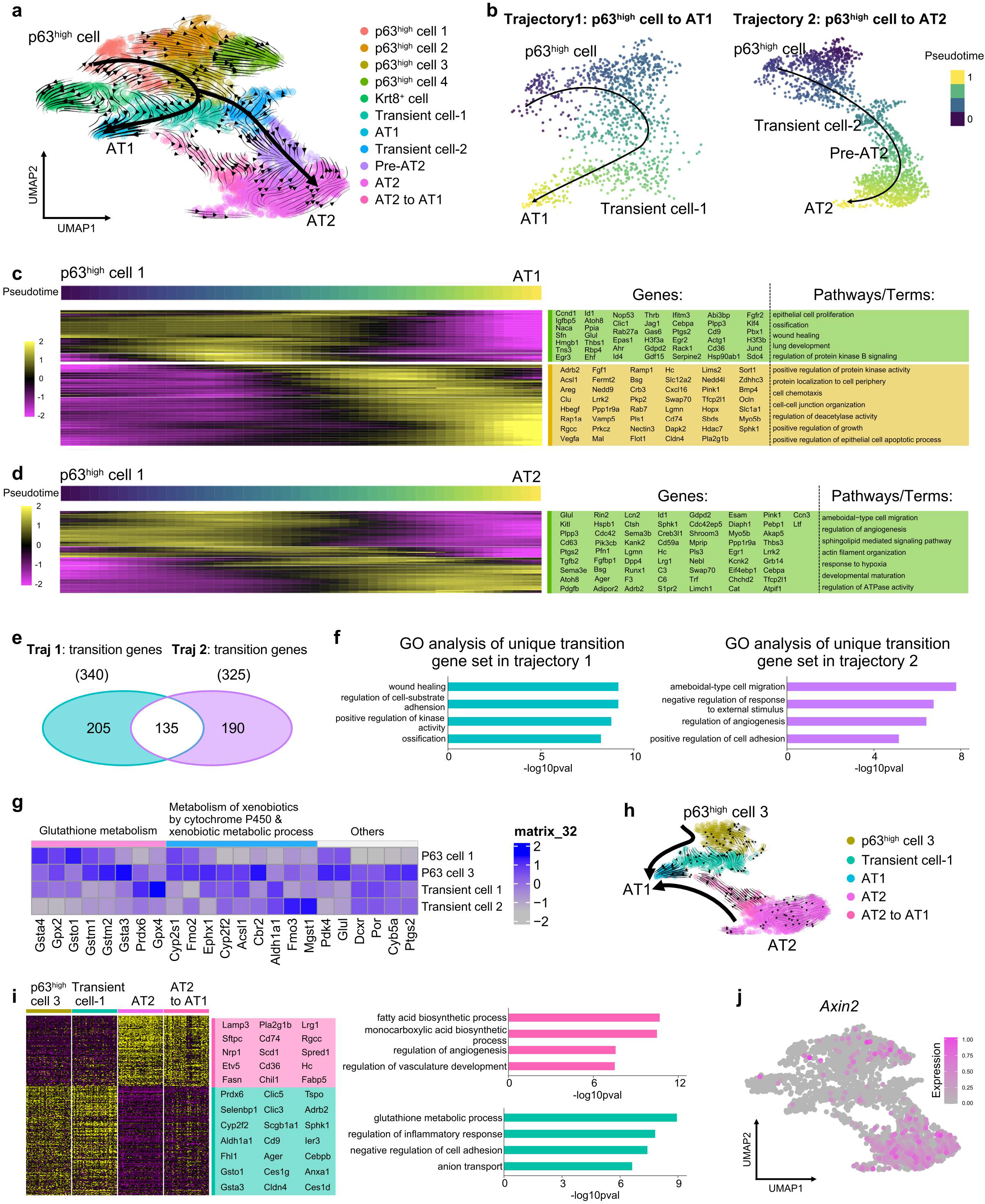
p63^+^ progenitors generate AT1 and AT2 cells through two distinct trajectories. **a**, RNA-velocity analysis of epithelial cells collected from bleomycin-treated lungs. Arrows indicate predicted lineage trajectories. **b**, Slingshot analysis of the pseudotimes of trajectory1 (p63^high^ cell to AT1 cell) and trajectory 2 (p63^high^ cell to AT2 cell) respectively. Arrows indicate the predicted lineage trajectories. **c-d**, The heatmap showing the gene expression patterns across the differentiation trajectory for transition genes. **e**, The Venn plot showing the common transition genes across two trajectories and unique transition genes for each trajectory respectively. **f**, Enrichment analysis showing that unique transition genes in each trajectory were enriched in the indicated pathways. **g**, Heatmap showing the levels of expression of genes associated with metabolism-related pathways across cell types as indicated. **h**, RNA-velocity analysis for AT1-related cell types as indicated. Arrows indicate the predicted lineage trajectories. **i**, Heatmap showing the differientiated expression genes of transient cell-1 and AT2-to-AT1 (left) and enriched pathways as indicated (right). **j**, UMAP plots showing *Axin2* expression in epithelial populations in bleomycin-treated lungs.

During the differentiation of p63^+^ progenitors, we found that p63-3 was a key node linking differentiation of p63-1 cells to AT1 or AT2 cells. By comparing the metabolism states of p63-1, p63-3, Transient 1, and Transient 2, we found that p63-3 cells have a glutathione metabolism and P450 associated metabolism signature, but the transient 1 or transient 2 displayed only one of them (Figure 3g). Additionally, we also found two clusters, transient cell-1 and AT2-to-AT1, that give rise to mature AT1 cells in the RNA velocity analysis (Figure 3h). These two transient clusters are distinct in transcriptome and maintain some of the signature genes of their cell-of-origin (Figure 3h,i). These data indicate that some p63^+^ progenitor cells may directly give rise to AT1 cells without first differentiating into AT2. Moreover, AT2 cells derived from p63^+^ progenitor cells express high levels of *Axin2*, a marker of Wnt-responsive alveolar epithelial progenitors^8,9^ (Figure 3i), indicating that a subset of newly generated AT2 cells from p63^+^ progenitors may possess stem cell capabilities. These data suggest that injury-induced p63^+^ progenitors give rise to AT1 and AT2 cells through two distinct routes, raising the possibility that p63^+^ progenitors are bipotent during alveolar epithelial regeneration.

### Sequential Genetic Tracing Reveals Conversion of Secretory to Alveolar Cells through a p63^+^ Progenitor Intermediate

Our lineage tracing data show that p63^+^ cells are induced after injury with bleomycin (Figure 1a-d) and do not arise from pre-existing p63^+^ cells (Extended Data Fig. 1d-f), suggesting a cell-of-origin distinct from the rare basal-like cells (p63^+^ LNEPs) that expand upon influenza. To identify the origins of injury-induced p63^+^ progenitors and determine their fates, we developed a dual recombinase-mediated genetic lineage tracing system to trace p63^+^ cells from specific parental lineages and follow their fate (Figure 4a). In this design, a lineage-specific CreER removes the loxP flanked Stop cassette in *p63-LSL-Dre* after Tam treatment, yielding a now-constitutive *p63-Dre* allele for recording p63^+^ cells thereafter. Using the *Rosa26-rox-Stop-rox-loxP-Stop-loxP-tdT (R26-Ai66*) reporter^28^, only cells that have expressed CreER and Dre are genetically labeled with tdT (Figure 4a), indicating the contribution of this specific cell lineage to p63^+^ cells. The genetic reporter tdT can then be used to examine whether p63^+^ cells contribute to alveolar cells during lung repair after injury. Thus, the advantage of this dual recombinase-mediated genetic system is that it enables simultaneous fate mapping of 3 sequential cell lineages in one mouse: cell of origin – p63^+^ cells – alveolar cells. We generated *Lineage-CreER;p63-LSL-Dre;R26-Ai66* mice (referred to “Lineage-p63 Tracer”) (Figure 4a). To verify whether the newly generated *p63-LSL-Dre* line faithfully monitors p63 gene activity, we crossed it with *ACTB-Cre* and use *R26-Ai66* to examine recombination in tissues (Extended Data Fig. 3a,b). We found that p63^+^ basal cells are labeled with tdT in *ACTB-p63 Tracer* mice, and we did not detect any tdT^+^ cells in *p63-LSL-Dre;R26-Ai66* mice (Extended Data Fig. 3c-f), demonstrating that the *p63-LSL-Dre* line can record p63 activity only after Cre-loxp recombination. We also crossed inducible *K5-2A-CreER* with *p63-LSL-Dre;R26-Ai66* mice and found that virtually all p63^+^K5^+^ basal epithelial cells were labeled in *K5-p63 Tracer* mice (Extended Data Fig. 3g-i).

**Figure 4.**
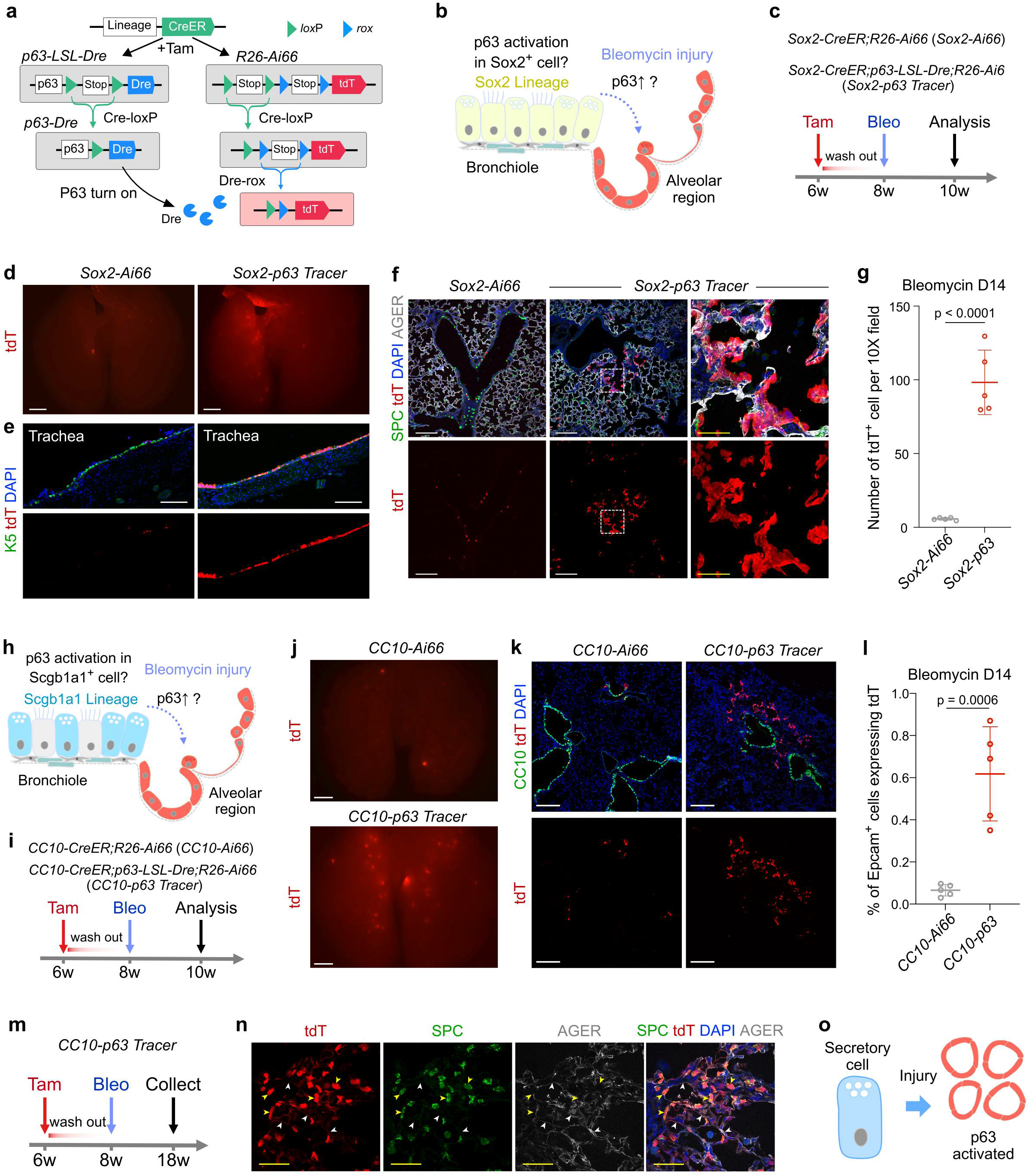
p63^+^ progenitors originate from secretory cells. **a**. A schematic illustrating the strategy for tracing both origin and fate of p63^+^ progenitors using dual recombinases. **b**, A cartoon image showing potential contribution of Sox2^+^ cells to p63^+^ progenitors. **c**, A schematic showing the experimental design. **d**, Whole-mount fluorescence images of *Sox2-Ai66* and *Sox2-p63* Tracer lungs. Scale bar, 2mm. **e**, Immunostaining for K5 and tdT on lung sections, Scale bar, 200 μm. **f**, Immunostaining for SPC, tdT, and AGER on lung sections. Scale bar, white, 200 μm; yellow, 50 μm. **g**. Quantification of the percentage of tdT^+^ cells. n = 5 mice for each group. **h**, A cartoon image showing potential contribution of secretory cells to p63^+^ progenitors. **i**, A schematic showing the experimental design of CC10-p63 system. **j**, Whole-mount images for tdT of bleomycin injured *CC10-p63 Tracer* or control lungs. Scale bar, 2 mm. **k**, Immunostaining for CC10 and tdT on lung sections. Scale bar, 200 μm. **l**, Quantification of the percentage of Epcam^+^ cells expressing tdT by FACS. **m**, A schematic showing the experimental design. **n**. Immunostaining for SPC, tdT, and AGER on lung sections collected at 2.5 months after injury. Scale bar, 50 μm. **o**, A cartoon image showing contribution of secretory cells to p63^+^ progenitors.

Given the proximity of p63^+^ cells to bronchioles (Figure 1d,f), we first explored the possibility that airway epithelial cells contribute to p63^+^ progenitor cells after injury. We crossed the airway epithelial cell-specific line *Sox2-CreER* with *p63-LSL-Dre;R26-Ai66* to generate *Sox2-p63 Tracer* mice, and examined if airway epithelial cells contribute to p63^+^ cells and their alveolar descendants (Figure 4b). We treated *Sox2-p63 Tracer* mice and littermate *Sox2-CreER;R26-Ai66* control mice with bleomycin at 2 weeks after Tam treatment, and analyzed lungs two weeks after injury (Figure 4c). As expected, a substantial number of p63^+^ basal cells in the trachea expressed tdT in *Sox2-p63 Tracer* mice, but not in *Sox2-CreER;R26-Ai66* mice (Figure 4d,e). In *Sox2-p63 Tracer* mouse lungs, we found that tdT^+^ cell patches were mainly located in alveolar regions near bronchioles (Figure 4f). Quantification showed that the number of tdT^+^ cells was significantly higher in lungs of *Sox2-p63 Tracer* mice than in *Sox2-CreER;R26-Ai66* control lungs (Figure 4g). Since the Sox2 lineage consists of multiple stem cell types including secretory, basal, and neuroendocrine cells, and considering the important role of secretory cells in distal airway regeneration both in mice and humans^15,20–22^, we next examined whether secretory cells contribute to p63^+^ progenitor cells after bleomycin injury. We generated *CC10-p63 Tracer (CC10-CreER;p63-LSL-Dre;R26-Ai66*) mice (Figure 4h,i). We found that tdT^+^ cell patches were located in peribronchiolar regions in *CC10-p63 Tracer* lungs, but not in *CC10-CreER;R26-Ai66* control lungs (Figure 4k,l). Using this dual genetic system, we observed that the CC10-derived p63^+^ cells further differentiate into SPC^+^ AT2 cells and AGER^+^ AT1 cells in *CC10-p63 Tracer* lungs collected at 10 weeks after injury (Figure 4m,n). Though it must be noted here that CC10 (*Scgb1a1*) transcript is somewhat promiscuous during injury, the above sequential genetic lineage tracing data strongly support a model wherein secretory cells contribute to p63^+^ progenitor cells that ultimately regenerate the alveolar epithelium after lung injury (Figure 4o).

Since AT2 cells are stem cells for alveolar regeneration^6,8,9^, we also examined whether AT2 cells convert to transient p63^+^ progenitors that subsequently generate new AT2 cells and AT1 cells. We crossed the AT2 cell-specific line *Spc-CreER* with *p63-LSL-Dre;R26-Ai66* to generate *Spc-p63 Tracer* mice (Figure 5a). We treated the mice with bleomycin 2 weeks after Tam treatment, and analyzed tissues at 2 weeks after injury (Figure 5b). We performed whole-mount lung imaging and immunostaining of sections and did not find a significant difference in the number of tdT^+^ cells in *Spc-p63 Tracer* and *Spc-CreER;Rosa26-Ai66* lungs (Figure 5c-e). As p63^+^ cells appeared in patches after injury (Figure 1e), the observation of scattered tdT^+^ cells similarly in both *Spc-p63 Tracer* and *Spc-CreER;R26-Ai66* control mice indicated that these tdT^+^ cells could be background signals from crosstalk by *Spc-CreER* with rox-Stop-rox (Cre-rox recombination). Nevertheless, tdT^+^ patches and tdT^+^ scattered cells could be distinguished in the lung, and quantification showed that minimal tdT^+^ patches were detected in both *Spc-p63 Tracer* and *Spc-CreER;R26-Ai66* control lungs (Figure 5d). To exclude the possibility that the lack of labeling was due to inefficient Cre-loxP recombination in AT2 cells, we isolated LysoTracker^+^ AT2 cells by FACS for genomic PCR analysis and found that Cre-loxP-mediated Stop was indeed removed in virtually all AT2 cells (Figure 5f), excluding the possibility of Cre-loxP recombination inefficiency.

**Figure 5.**
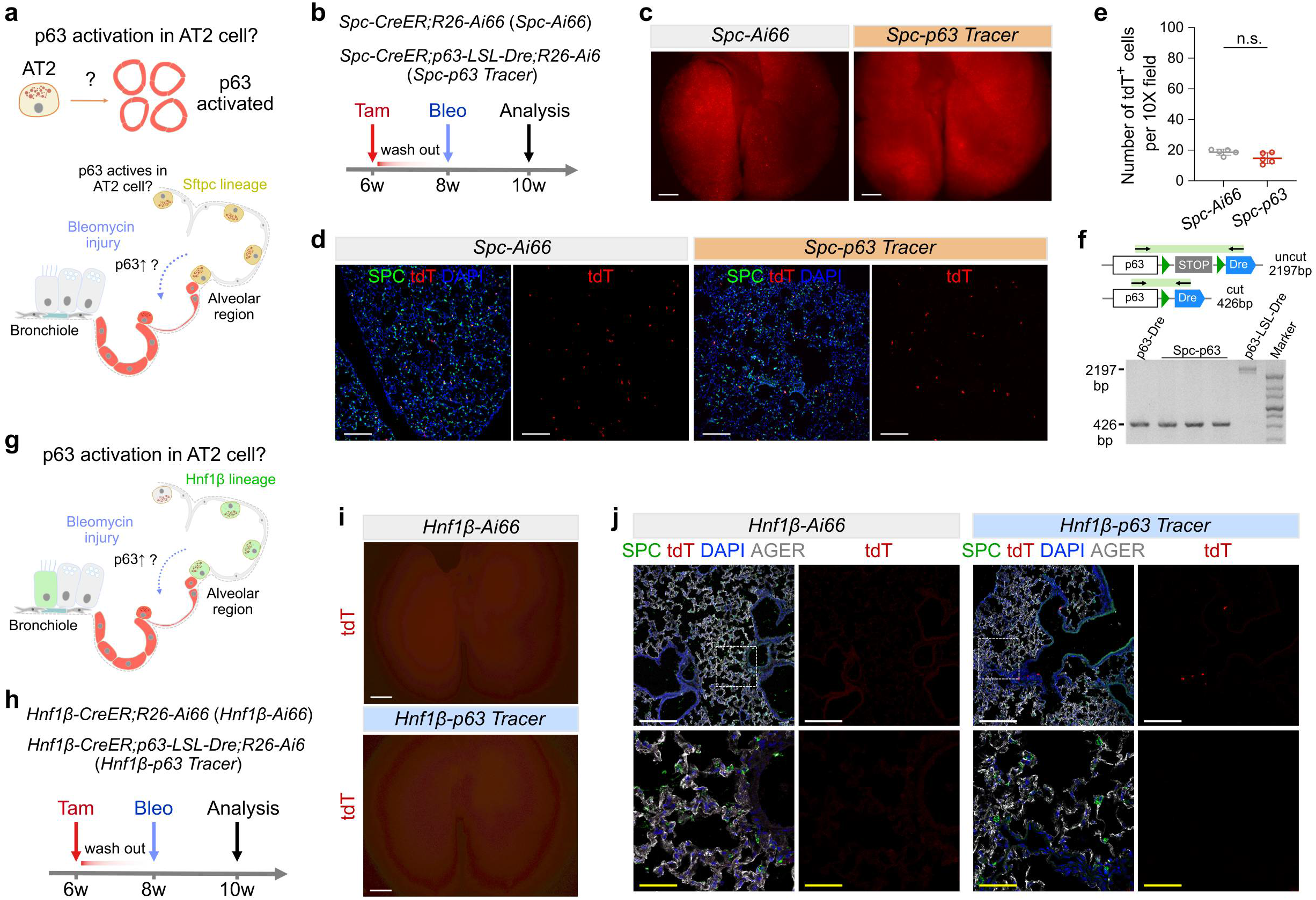
AT2 and multi-ciliated cells minimally contribute to p63^+^ progenitors. **a,b**, A schematic showing the experimental design. **c**, Whole-mount fluorescence images of *Spc-Ai66* and *Spc-p63 Tracer* lungs. Scale bar, 2 mm. **d**, Immunostaining for SPC and tdT on lung sections. Scale bar, 200 μm. **e**, Quantification of tdT^+^ cells per 10x field. n = 5 mice for each group. **f**, PCR primer design for detection of the recombination efficiency of LSL-Dre by *Spc-CreER* line. **g**,**h**, A schematic showing the experimental design. **i**, Whole-mount fluorescence images of *Hnf1β-Ai66* and *Hnf1β-p63 Tracer* lungs. Scale bar, 2 mm. **j**, Immunostaining for SPC, tdT, and AGER on lung sections. Scale bar, 100 μm.

Due to the background Cre-rox crosstalk signals in *Spc-p63 Tracer*, we next generated a new AT2 cell-specific Cre driver *Hnf1β-CreER* mice (Extended Data Fig. 4a). We found *Hnf1β-CreER* specifically and efficiently labels SPC^+^ AT2 cells but not AGER^+^ AT1 cells (Extended Data Fig. 4b-d). Additionally, *Hnf1β-CreER* also targets Acy-T^+^ ciliated cells but not CC10^+^ club cells (Extended Data Fig. 4e). We crossed *Hnf1β-CreER* with *p63-LSL-Dre;R26-Ai66* to generate *Hnf1β-p63 Tracer* mice (Figure 5g). We treated *Hnf1β-p63 Tracer* and *Hnf1 β-CreER;R26-Ai66* mice with bleomycin at 2 weeks after Tam, and collected tissues at 2 weeks after injury (Figure 5h). Whole-mount fluorescence imaging and sectional immunostaining revealed sparse tdT^+^ cells and virtually no tdT^+^ patches in *Hnf1β-p63 Tracer* mice (Figure 5i,j). We did not detect any tdT^+^ cells in *Hnf1β-CreER;R26-Ai66* mice lungs, demonstrating no Cre-rox crosstalk contamination by this system. The above data demonstrate that AT2 cells and ciliated cells in airways contribute minimally, if at all, to p63^+^ progenitors that emerge after injury.

### p63 is Required for Efficient Alveolar Regeneration from Secretory Cells

We next asked whether p63 activation during injury is functionally required for generation of p63^+^ progenitors (Figure 6a). To simultaneously trace secretory cell-derived p63^+^ progenitors and delete the p63 gene specifically in CC10^+^ secretory cells, we generated a conditional flox allele for p63 (*p63-f1*), in which exon 5 to exon 7 are flanked by loxP sites (Figure 6b). We first validated this new *p63-fl* line by crossing it with an inducible *K5-2A-CreER* driver, which specifically and efficiently deleted the p63 gene in basal cells (Extended Data Fig. 5a,b). We then crossed the *p63-fl* line with *CC10-p63 Tracer* mice to generate *CC10-CreER;p63-LSL-Dre/flox;R26-Ai66* mice (*CC10-p63 Tracer;p63-fl*), in which one p63 allele was null and another p63 allele was conditional flox (Figure 6b). Experimental data showed that *p63-LSL-Dre* was a null allele, as homozygous mice for LDL-Dre had no p63 expression and exhibited the same defects in limb, craniofacial and epithelial development as p63^−/−^ mice (Extended Data Fig. 5c), consistent with previous studies^29,30^. We treated *CC10-p63 Tracer;p63-fl* mice with Tam, induced bleomycin injury after 3 weeks, and collected lungs at 2 weeks after injury (Figure 6b). *CC10-p63 Tracer* was used as control, as one p63 allele remained intact (Figure 6b). PCR analysis showed around 70% knockout efficiency in secretory cells (Figure 6c). Immunostaining on tissue sections revealed a significant reduction of tdT^+^ cells in *CC10-p63 Tracer;p63-fl* compared with *CC10-p63 Tracer* lungs (Figure 6d,e). Of note, we could hardly detect tdT^+^ patches in *CC10-p63 Tracer;p63-fl* compared with *CC10-p63 Tracer* lungs, indicating that p63 expression was required for expansion and/or survival of p63^+^ progenitors during injury.

**Figure 6.**
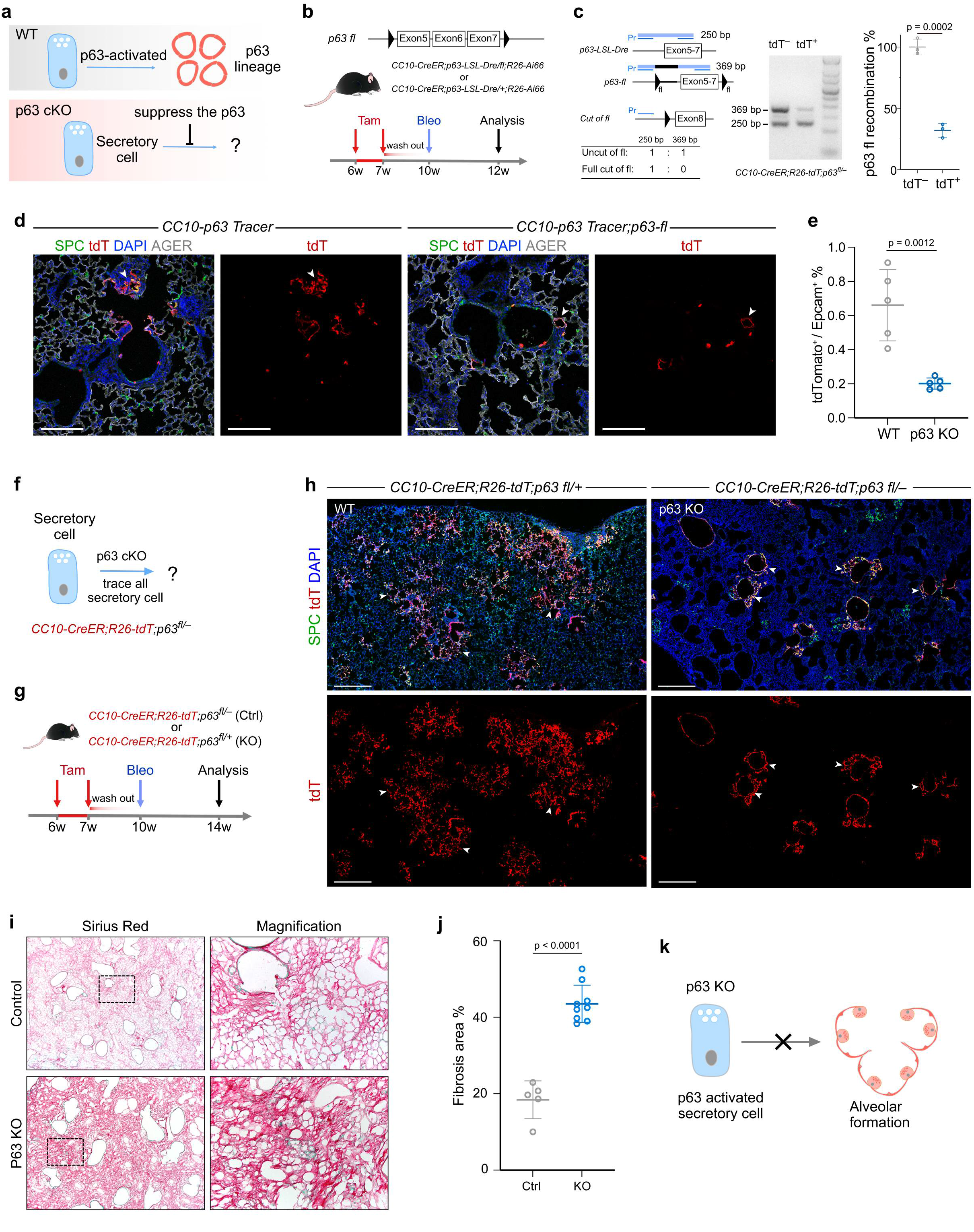
p63 is required for contribution of secretory cells to alveolar regeneration. **a**, A schematic showing the experimental design. **b**, A schematic showing conditional p63 knockout in secretory cells and simultaneous tracing of p63^+^ progenitors. **c**. PCR detection of the ratio of *p63-LSL-Dre* band and *p63-fl* band infers the recombination efficiency of *p63-f1* by *CC10-CreER* line (left lane). Right panel shows ~70% in recombination efficiency. **d**, Immunostaining for SPC, tdT, and AGER on lung sections. Arrowheads indicate tdT^+^ alveolar cells. Scale bar, 100 μm. **e**, Quantification of the percentage of Epcam^+^ cells expressing tdT by FACS. n = 5 mice for each group. **f**, A cartoon image showing tracing p63-depleted secretory cells. **g**, A schematic showing the experiment design. **h**, Immunostaining for SPC and tdT on lung sections. Arrowhead indicate tdT^+^ alveoli. Scale bar, 500 μm. **i**, Sirius-red staining of p63 KO and control mice. **j**, Quantification of fibrosis area in **i**. n = 5 mice in control and 9 mice in p63 KO mice. **k**, A cartoon image showing p63 KO impairs alveolar generation from secretory cells.

As our data suggested that airway secretory cells contributed to alveolar regeneration through p63^+^ progenitors, we independently addressed whether p63 activation was required for this lineage conversion by tracing p63-depleted secretory cells. We generated *CC10-CreER;R26-tdT;p63^f1/-^* mutant mice and littermate *CC10-CreER;R26-tdT;p63^f1/-^* control mice to trace secretory cells with p63 deleted and a functional copy of p63, respectively. We treated mice with bleomycin at 3 weeks after Tam administration and collected lungs for analysis at 4 weeks after injury (Figure 6g). Immunostaining for tdT and SPC on injured lung sections revealed that a substantial number of tdT^+^ patches in alveolar regions adopt alveolar epithelial cell fates in *CC10-CreER;R26-tdT;p63^f1/-^* control mice but fewer in *CC10-CreER;R26-tdT;p63^f1/-^* mutant mice (Figure 6h). Sirius Red staining revealed more severe fibrosis in *CC10-CreER;R26-tdT;p63^f1/-^* mutant lungs than in *CC10-CreER;R26-tdT;p63^f1/-^* controls (Figure 6i,j), indicating impaired alveolar regeneration in lungs with p63 deficient secretory cells. Taken together, these data suggest that p63 activation is functionally required for efficient contribution of secretory cells to alveolar regeneration after bleomycin-induced lung injury.

## Discussion

In this study, we identified an injury-induced p63^+^ progenitor that contributes to alveolar regeneration and by applying dual recombinase-mediated genetic lineage tracing, we elucidated sequential conversion of secretory cells to p63^+^ progenitors and then alveolar cells during repair after injury (Figure 7). These transitional p63^+^ progenitors appear to contribute to alveolar cells through two distinct routes during their differentiation into AT1 and AT2, respectively. We also found that p63, a key transcription factor responsible for maintaining basal cell identity, is functionally required for the contribution of secretory cells to alveolar epithelial cells after lung injury (Figure 7). The identification of transient p63^+^ progenitors for alveolar regeneration after injury improves our understanding of epithelial cell fate plasticity and dynamic cell states during cell lineage conversion. Further exploration of the molecular signals that control injury-induced progenitors would provide us with a potential therapeutic avenue for lung epithelial cell regeneration.

**Figure 7.**
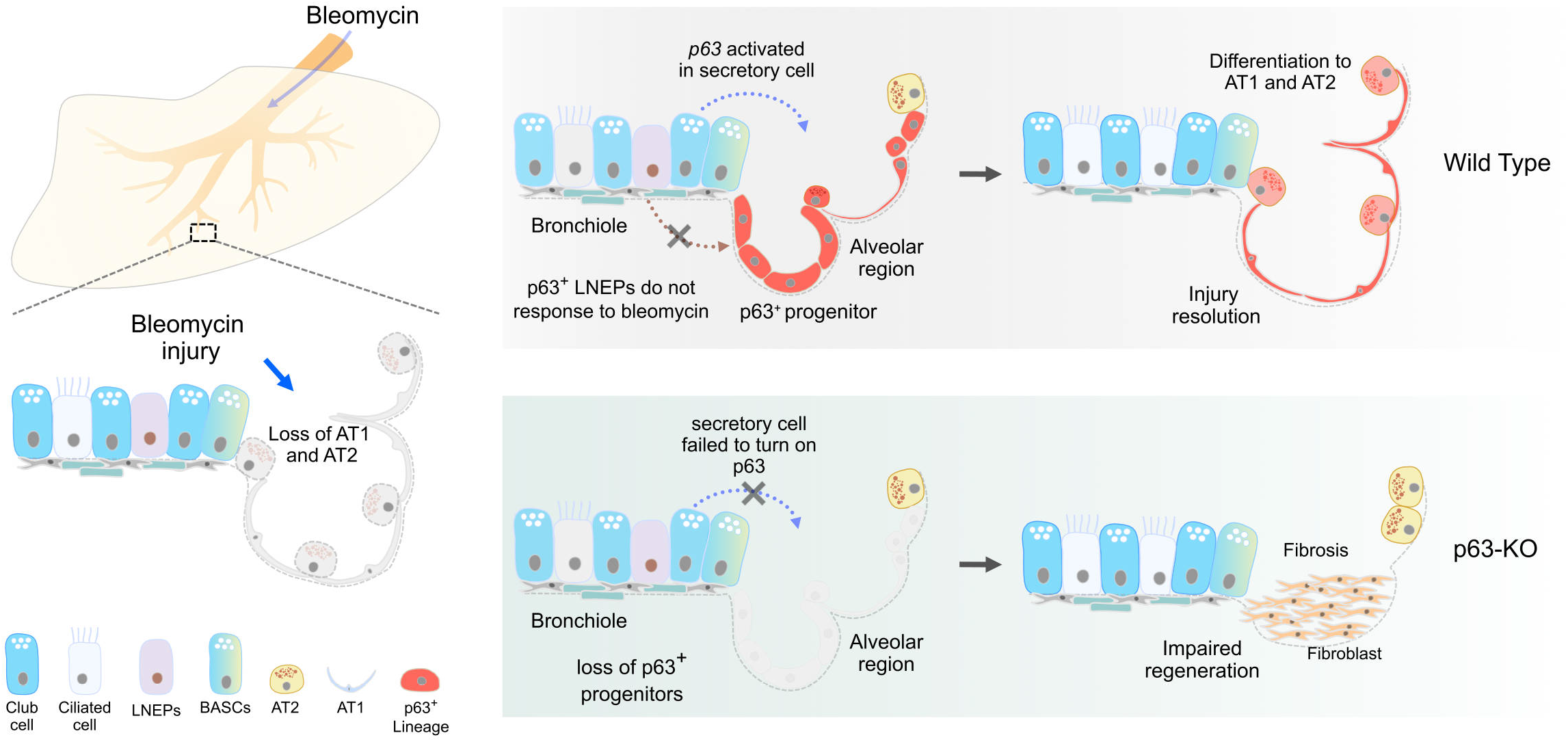
Injury-induced p63^+^ progenitors regenerate alveoli. A cartoon image showing contribution of secretory cells to injury-induced p63^+^ progenitors that regenerate alveolar cells. p63 deletion significantly impairs alveolar regeneration and leads to severe fibrosis.

To understand the source and fate of p63^+^ progenitors during injury, we first used *p63-CreER* to label pre-existing p63^+^ basal cells and intrapulmonary p63^+^ progenitors before injury^25,31^. After bleomycin injury, we found these pre-labeled p63^+^ cells do not contribute appreciably to p63^+^ cells in damaged lung parenchyma, which is in stark contrast to multiple reports demonstrating that essentially all of the p63^+^ Krt5^+^ expansion after influenza can be attributed to pre-existing intrapulmonary p63^+^ progenitors^16,18,25^. Instead, the honeycomb-like p63^+^ cells appearing after bleomycin injury were *de novo* generated by p63^−^ cells. The injury-induced p63^+^ progenitors expanded significantly and contributed to alveolar cells. To directly link the origin of p63^+^ progenitors and their ultimate alveolar epithelial cell fate, we developed a sequential genetic tracing strategy that clearly demonstrated that secretory cells expressing CC10 contribute to p63^+^ cells that subsequently differentiate into alveolar epithelial cells. Recent works on the human distal airway have identified new cell populations AT0s and RASCs, which are intermediate stages between secretory cells and AT2s, in homeostasis and disease^20,21^. The existence of these intermediate cell populations in humans suggests the regeneration program of the human distal epithelial cell may involve the transdifferentiation of secretory cells to alveolar epithelial cell lineage, which our studies here suggest involves transient upregulation of p63. These results provide important clarifications about the nature of injury-induced expansion of p63^+^ progenitor cells during lung injury. Specifically, the relative lack of alveolar differentiation capacity of influenza-induced p63^+^ Krt5^+^ cells appear to reflect their basal cell-like origin, whereas the p63^+^ cells induced by bleomycin injury, arising from secretory cells, seem to “remember” their more distal lung origin and are capable of contributing to euplastic alveolar regeneration.

The injury-induced p63^+^ progenitors adopt two new transient cell stages before differentiating into AT1 and AT2. The Transient stage 1 gradually differentiates to AT1 and the Transient stage2 differentiates to Pre-AT2 and further converts to mature AT2. During the differentiation, we identified an important cell state node, p63-3, which further differentiated to Transient 1 or Transient 2. The p63-3 shares features with p63^+^ progenitors including high proliferation potential and a similar metabolism program, and also expresses some metabolism genes that are highly expressed in Transient 1 and Transient 2. These intermediate stages indicated the p63^+^ progenitor is a bipotent progenitor which gives rise to AT1 and AT2. The detection of Transient 1 state suggests the p63-activated progenitor can directly differentiate to AT1 without differentiating to AT2. Recent studies reported that during recovery after lung injury, AT2 differentiate into AT1 through a K8^+^ intermediate stage named DATP (damage-associated transient progenitors) or transient stage called PATS (pre-alveolar type-1 transitional cell state)^26,27,32^. These transitional states correlate with abnormal epithelial cells associated with defective fibrotic foci in human lung diseases^32^. Our data showed the p63^+^ progenitor also adapts to this transient stage, with a similar transcription signature, suggesting the possibility that the p63^+^ progenitor may also respond to the inflammation stimulation and be activated by IL-1β like AT2. However, our RNA velocity analysis showed that K8^+^ transient cells are in a terminal state and their differentiation to mature AT1 is not obvious. These data suggest that transdifferentiation of p63^+^ progenitors to mature AT1 may not involve a K8^+^ intermediate. The detailed characterization of p63^+^ progenitors and their descendants offer new insights into how progenitors react in response to injury and generate different alveolar cells through distinct programs.

p63 plays an important role in the development of stratified epithelial stem cells and the maintenance of the stem cell pool in homeostasis^31,33,34^. p63 is activated in epithelial cells in response to injury, and p63 is a crucial transcription factor for regulating key genes in airway basal cells^35^. p63 regulates the expression of adhesion-associated genes and cell survival^36^, which are also detected in p63^+^ progenitors. Unlike “true” basal cells, the injury-activated p63 in secretory cell lineages and these p63^+^ progenitors did not robustly express basal cell markers. The reason for the directional differentiation of injury-induced p63^+^ progenitors to alveolar epithelial cell lineages rather than basal lineages remains unclear, but we posit that this differential plasticity is reflective of both the differential cells of origin as well as the different natures of the inflammatory environment between sterile and infectious injuries. One possible explanation is that p63^+^ progenitors gradually lose the p63 activation and turns to Transient 1 and Transient 2 stages when they migrate into an environment that favors alveolar differentiation. It could also be explained by the intrinsic heterogeneity of the secretory cells, as the proximal secretory cell can dedifferentiate and reform basal stem cells when their parental basal stem cells are ablated^37^, while distal secretory cells convert to p63^+^ progenitors that preferentially differentiate into alveolar cells. Intriguingly, we demonstrate that adoption of a p63^+^ intermediate after bleomycin injury seems to empower club cells to participate in repair, whereas loss of p63 is also required for p63^+^ Krt5^+^ cells to appreciably contribute to alveolar repair after influenza^18^, reinforcing the notion that dynamics of p63 expression are critical for both injury responses but also the resolution of epithelial remodeling. However, what drives some secretory cells to activate the expression of p63 and their translocation into the injured lung parenchyma remains unclear. Further studies are needed to clarify the regulation pathway of secretory cell activation, migration, and their subsequent fate conversion, which may offer new insight in our understanding stem cell plasticity and regenerative capability after lung injury.

## Acknowledgments

This work was supported by the National Key Research & Development Program of China (2019YFA0110403, 2019YFA0802000 to B.Z., 2022YFA104200 to Y.L.), the CAS Project for Young Scientists in Basic Research (YSBR-012 to B.Z.), the Youth Innovation Promotion Association CAS, the Shanghai Pilot Program for Basic Research – CAS, Shanghai Branch (JCYJ-SHFY-2021-0 to B.Z.), the National Science Foundation of China (32050087, 82088101, 32100592, 32170848, 32100585, 32100648 to B.Z., W.P., Y.L., H.Z., and K.L.), the Program for Guangdong Introduction Innovative and Entrepreneurial Teams (2017ZT07S347 to B.Z.), the support from the XPLORER PRIZE and New Cornerstone Science Foundation (to B.Z.), and the support from Boehringer-Ingelheim (to B.Z.), and by Shanghai Municipal Science and Technology Major Project (to B.Z.). We thank D. Huang and H.Y. Wu for acquiring light-sheet imaging data. We also thank the Shanghai Model Organisms Center, Inc. (SMOC) for mouse generation; and institutional animal facilities for mouse husbandry.

## Author Contributions

Z.L. and B.Z. conceived and designed the project. Z.L. performed single-cell RNA-sequencing experiments and analysis. K.L., W.P., Y.L., and H.Z. bred mice, performed mouse experiments, and analyzed the data; Y.X., A.E.V., and A.G. provided intellectual input and revised the manuscript; B.Z. interpreted the data and drafted the manuscript.

## Competing Interests

Authors declare no competing interests.

## Online Methods

### Mice

All mice used in experiments were housed and bred in accordance with the guidelines of the Institutional Animal Care and Use Committee (IACUC) of the Center for Excellence in Molecular Cell Science, Shanghai Institutes of Biological Sciences, Chinese Academy of Science. The *R26-tdTomato* (*R26-tdT*)^24^, R26-Ai66^28^, ACTB-Cre^38^, Sox2-CreER^39^, *CC10-CreER^40^* and *Spc-CreER^41^* mouse lines were reported previously. *p63-CreER* and *p63-fl* mouse lines were generated by homologous recombination using ES cells as previously described^42^. *K5-2A-CreER, p63-LSL-Dre, Hnf1β-2A-CreER*, and *p63-DreER (p63^−/+^*) mouse lines were generated by homologous recombination using CRISPR/Cas9 as previously described^43^. In brief, the *p63-CreER* line was generated by inserting the P2A-CreER^T2^-WPRE-PolyA sequence at the end of exon2 in the *Trp63* gene. The same P2A-CreERT2-WPRE-PolyA sequence was inserted at the stop codon of the *Krt5* gene or *Hnf1β* gene to generate *K5-2A-CreER* and *Hnf1β-2A-CreER* mouse lines. For the *p63-LSL-Dre* line, the cDNA encoding loxP-stop-loxP-Dre-WPRE-PolyA was inserted into the ATG of the *Trp63* gene. For the *p63-DreER* line, the cDNA sequence of DreERT2-WPRE-PolyA was inserted into the start site of exon4 of the *Trp63* gene as previously described^44^. The *p63-fl* mouse line was generated by inserting two loxP sites flanking the sequence of exon 5 to exon 7 of the *Trp63* gene. The *p63-fl, K5-2A-CreER, p63-LSL-Dre, Hnf1β-2A-CreER*, and *p63-DreER* mice were generated by Shanghai Model Organisms Center, Inc. (SMOC), and the *p63-CreER* mouse was generated by GemPharmatech Co, Ltd. All mice involved in this study were maintained on C57BL/6/129 background and kept under a 12-hour day and light cycle. To lineage label p63^+^ progenitors, the *p63-CreER* mice was crossed with *R26-tdT* mice to generate *p63-CreER;R26-tdT*. For *p63-Tracer* mice, the *Sox2-CreER, CC10-CreER, Spc-CreER*, or *Hnf1β-2A-CreER* mice were crossed with *p63-LSL-Dre;R26-Ai66* mice, respectively, to determine the origin of p63^+^ progenitors that emerge after injury. The *p63-fl* strain was crossed with *CC10-p63 Tracer* mice to generate a conditional knockout of p63 in secretory cells. Adult mice received 0.2 mg tamoxifen (TAM) per gram of mouse body weight by oral gavage for indicated doses and at indicated times. Both male and female mice were included in the study.

### Genomic PCR

Genomic DNA was extracted from the mouse tail or embryonic yolk sac as previously described^42^. Mouse tissues were treated with 100 μg ml^−1^ Proteinase K in lysis buffer (100 mM Tris HCl, pH7.8, 5mM EDTA, 0.2% SDS, and 200 mM NaCl) overnight at 55°C, followed by centrifugation at 25,000 g for 8 minutes. DNA was precipitated in isopropanol, washed by 70% ethanol, and dissolved in distilled water. All mice were genotyped with specific PCR primers that distinguish knock-in allele from wild-type allele. Sequences of all primers were included in Supplementary Table1.

### Bleomycin injury

Bleomycin was delivered by intra-tracheal instillation as described previously^45^. In brief, mice were anesthetized by intra-peritoneal (i.p.) injection of 2,2,2-Tribromoethanol (200mg kg^−1^, Sigma-Aldrich, T48402), and the body was stably fixed on a board. The tribromoethanol was dissolved in tert-Amyl alcohol (GCS, 40090360) to 1.6 mg ml^−1^ and then dissolved to 20 mg ml^−1^ in PBS. The 1 U kg^−1^ dose of bleomycin (Sigma, B8416) dissolved in sterile PBS (Invitrogen, 10010049) was administrated via a cannula needle that plugged in the trachea and inhaled with the breath. The control group mice inhaled the same corresponding volume of sterile PBS. The lung tissues were collected at different times after injury as indicated in each experiment.

### Lung tissue collection and whole-mount fluorescence microscopy

Lung perfusion was performed as previously described^45^. After mice were euthanized, the blood in the lung was flushed out by perfusion with 10 to 15 ml cold PBS through right ventricle. The lung was perfused with 1.5 ml 4% paraformaldehyde (PFA) through the trachea and the collected tissue was fixed in PFA for 1 h at 4°C, and then washed in PBS for three times. The fixed tissues were placed stably on 1% agar gel for whole-mount imaging using a Zeiss stereo microscope (AxioZoom V16). For magnification of specific regions, we used the automated z-stack images acquired with a Zeiss stereoscope (AxioZoom V16) as previously described^11^.

### Tissue clearing and lightsheet microscopy

The lung was cleared by the advanced CUBIC method as reported^46^. Mice were anesthetized by i.p. injection of tribromoethanol (200mg kg^−1^, Sigma-Aldrich, T48402). The lung was perfused by injecting 10 ml cold PBS through the right ventricle and then the lung was perfused with and immersed in 4% PFA at 4 °C for 1h. The fixed tissue was washed in 10 ml PBS twice at room temperature for 4 hours. The washed tissue was then immersed in 10 ml DAPI (1:1000) containing 1/2 water-diluted-reagent-1 with shaking 60 r.p.m. on orbital shaker at 37 °C for 6 h. The reagent-1 is a mixture of 25 wt% of urea, 25 wt% of Quadrol, 15 wt% of Triton X-100 and 35 wt% of dH_2_O. Then, the clearing solvent was switched to 10 ml DAPI containing reagent-1 and the sample was gently rotated at 37 °C overnight. The reagent-1 was replaced every day and the tissue was cleared by ~7 days after treated with reagent-1. The cleared tissue was immersed in reagent-1 at room temperature avoiding from the light for short term preservation and prepared for imaging steps. Images was taken by LiTone LBS Light-sheet Microscope (Light Innovation Technology Limited) with Olympus XLFLUOR4X/340 objective lens (N.A. = 0.28) and further image processing was performed by Imaris (Oxford Instruments, 9.2.4).

### Immunostaining

Immunostaining on tissue sections were performed according to protocols described previously^47^. All tissues were treated with 4% PFA for 1 hour at 4 °C and washed three times in PBS, and dehydrated in 30% sucrose over night at 4 °C and embedded in OCT (Sakura). For staining, the cryosections were washed in PBS, incubated in blocking buffer (5% normal donkey serum from Jackson Immunoresearch, 0.1% Triton X-100 in PBS) for 30 minutes at room temperature, followed by incubation with the primary antibodies overnight at 4°C. The following antibodies were included in this work with company names, catalog number, and indicated dilution: p63 (Abcam, ab735, 1:200), Ki67 (Abcam, ab15580, 1:200), SPC (Millipore, AB3786, 1:200), CC10 (Abcam, ab213202, 1:300), Acetylated-tubulin (Sigma, T7451, 1:500), AGER (R&D systems, MAB1179, 1:200), Aqp3 (Invitrogen, PA578811, 1:500), WFDC1 (Invitrogen, PA5-116731, 1:100), Scgb3a2 (R&D systems, AF3545, 1:500), Krt8 (DSHB, Troma-I-C, 1:500), tdTomato (tdT, Rockland, 600-401-379, 1:500; or Rockland, 200-101-379, 1:500). Signals were developed with Alexa fluorescence antibodies (Invitrogen, 1:500). Nuclei were counterstained with 4’6-diamidino-2-phenylindole (DAPI, Vector lab). For the staining of P63, additional antigen retrieval by incubating the section in antigen retrieval buffer (10mM Sodium Citrate, 0.05% Tween 20, pH 6.0 or 1X of Beyotime, P0088) at around 98 °C for 10 minutes was needed before the donkey serum blocking procedure. Immunostaining images were acquired by Zeiss confocal microscope (LSM880) or Nikon A1 confocal microscope. Imaging of all immunostained slides for side-by-side comparisons was performed under the same exposure and contrast conditions using the same confocal microscope.

### Lung epithelial cell isolation

The lung epithelial cells isolation procedure was same as previously described^11^. The whole process included two rounds of digestion by two different enzyme cocktails. Enzyme mixture 1 included collagenase type I (500 U ml^−1^, Gibco, 17100–017), elastase (4 U ml^−1^, Worthington Biochemical Corporation, LS002279), and dispase (5 U ml^−1^, BD Biosciences, 354235) in DMEM/F12 (Gibco, 10565018). Enzyme mixture 2 contained 0.1% Trypsin-EDTA (Gibco, 25200072) and DNase I (0.33 U ml^−1^; Worthington Biochemical Corporation LS002139) in DMEM/F12 (Gibco, 10565018). In brief, mice were euthanized by CO_2_ inhalation and their lungs were perfused with 10 ml cold PBS through the right ventricle. After injecting 1.5 ml enzyme mixture 1 through trachea, the lung lobes were collected and cut in to pieces, and additional 2.5 ml enzyme mixture 1 was added before incubating the sample at 37 °C for 20 minutes on orbital shaker (~100 r.p.m). The digestion was ended by adding equal volume of cold DMEM/F12 with 10% of FBS (Gibco, 10099141) to the digestion medium. The tissue was disturbed by pipetting before centrifuged with 1000g at 4 °C for 5 minutes. The supernatant was discarded and the pellet was resuspended with 2 ml enzyme mixture 2 for incubating at 37 °C for 20 minutes. The digestion was stopped by adding equal volume of DMEM/F12 containing of 10% FBS and then filtered with 100 μm strainer and centrifuged with 1000g at 4 °C for 5 minutes. The pellet was resuspended by 2-3 ml red blood cell lysis buffer (eBioscience, 00-4333-57) and incubated at room temperature for 3 minutes and then was stopped by adding 3 ml cold PBS. The sample was filtered through 40 μm strainer and then centrifuged and resuspended in PBS. For MACS and FACS combined isolation, the cells was incubated with Epcam-APC (1:100, eBioscience, 17-5791-82) at 4 °C for 30 minutes and ended by adding cold PBS and centrifuged with 1000g at4 °C for 5 minutes. For each adult lung, the pellet was resuspended in 80 μl of PBS and followed by 320 μl of anti-APC MicroBeads (Miltenyi Biotec, 130-090-855), then the cells and beads were mixed well by pipetting, followed by incubation at 4 °C for 15 minutes. After washing the sample with PBS and centrifuging with 1000g at 4 °C for 5 minutes, the pellet was resuspended in 500 ul of PBS and performed MACS positive isolation by autoMACS Pro Separator (Miltenyi Biotec). The positive selected cells were stained with DAPI and were used for FACS isolation by gating DAPI^−^ Epcam^+^ tdTomato^+^ cells. For isolation of *CC10-CreER* labelled secretory cells, the staining process included extra antibodies, CD45-PE-Cy7 (1:400, eBioscience, 25-0451-82), and the gating strategy was changed to DAPI_-_ CD45^−^ Epcam^+^ tdTomato^+^ cells or DAPI^−^ CD45^−^ Epcam^+^ tdTomato^+^ cells.

For isolation of AT2 cells, a procedure for staining with LysoTracker-DND-26 (Invitrogen, L7526) at 37 °C for 15 minutes was needed after the antibody staining process. The DAPI^−^ CD45^−^ Epcam^+^ LysoTracker^+^ cells were used for AT2 isolation gating strategy.

### Single-cell RNA Sequencing

The suspended cells were loaded on a 10x Chromium controller to generate GEMs, which were further processed into single cell 5’ gene expression libraries using the Chromium Single Cell 5’ Library Kit (v2 Chemistry). Sequencing was performed on Illumina NovaSeq 6000 PE150 platform. Raw fastq files first were cleaned by Trim Galore with parameter “-q 20–phred33–stringency 3–length 20 -e 0.1”. Each scRNA-seq data set was processed by CellRanger (v6.1.1) pipeline to perform the alignment against the modified mouse reference genome mm10, filter the alignment and count barcodes and UMIs. First, quality control was performed by R package seurat^48^ (v4.1.2). Cells with a mitochondrial-fraction above 0.1 or 0.08 and less than 2500 genes were removed from the analysis. Next, different time point data sets were integrated by MNN algorithm^49^ wrapped in R package batchelor (v1.10.0). The R package DBSCAN^50^ (v1.1-10) was used to identify outlier cells which were subsequently removed from further analysis. The downstream analysis was mainly performed using the R package seurat (v4.1.2) with customized parameters. GO enrichment analysis was performed by R package clusterProfiler^51^ (v4.2.2) or metascape (https://metascape.org/gp/index.html#/main/step1). We employed scvelo^52^ (https://github.com/theislab/scvelo) to infer the future state of individual cells through information of unspliced and spliced transcripts. Selected gene set signature activity analysis were performed by R package AUCell^53^ (v1.19.1). Pseudotemporal ordering along the trajectory of the cellular embeddings were inferred with the R package slingshot^54^ (v2.2.1). Differentially expressed genes along the trajectories were identified using the R package tradeSeq^55^ (v1.8.0). Gene expression patterns along the trajectories were visualized using the R loess() function with default parameters.

### Sirius Red staining

The sections were immersed in PBS for 5 minutes to wash out the OCT in the tissues and then immersed in Bouins’ solution for 12-24 hours. Tap water was used to wash out the Bouins’ solution on sections till the yellow color disappeared, followed by staining in Fast green for 3 minutes and subsequent 3 times of washing by tap water. Then the sections were soaked in 1% acetic acid for 1 minute and washed with ddH_2_O before they were embedded in 0.1% Sirius red for 1 to 1.5 min. The stained sections were washed by ddH_2_O and dehydrated with 95% ethanol, 100% ethonol, and Xylene sequentially. The well dehydrated sections were mounted by Permount™ Mounting Medium (Shanghai Maokang Biotechnology, MM1411-100ML). All processes were performed at room temperature.

### Statistics

Tissue samples were randomized and blinded for analyzers and for each experiment no less than 5 biological samples for each group were collected. Data were presented as means ± SD. To-tailed unpaired Student’s *t* test was used for comparison of differences between two groups, and ANOVA followed by Tukey’s method was used for multiple comparisons in statistical analysis. *P* < 0.05 was considered to be statistically significant. The biological number n in each figure indicates mouse number.

## Data availability

All of the data generated or analyzed during this study are included in Figs. 1–7, Extended Data Figs. 1–5, and Supplementary Tables 1. scRNA-seq data that support this study have been deposited in the Gene Expression Omnibus (GEO) of NCBI (Accession number GSE225187).

## Notes

### Competing Interest Statement

The authors have declared no competing interest.

