## Supplementary Figures 1-5 and Table 1 for "Airway secretory cell-derived p63^+^ progenitors contribute to alveolar regeneration after sterile lung injury"

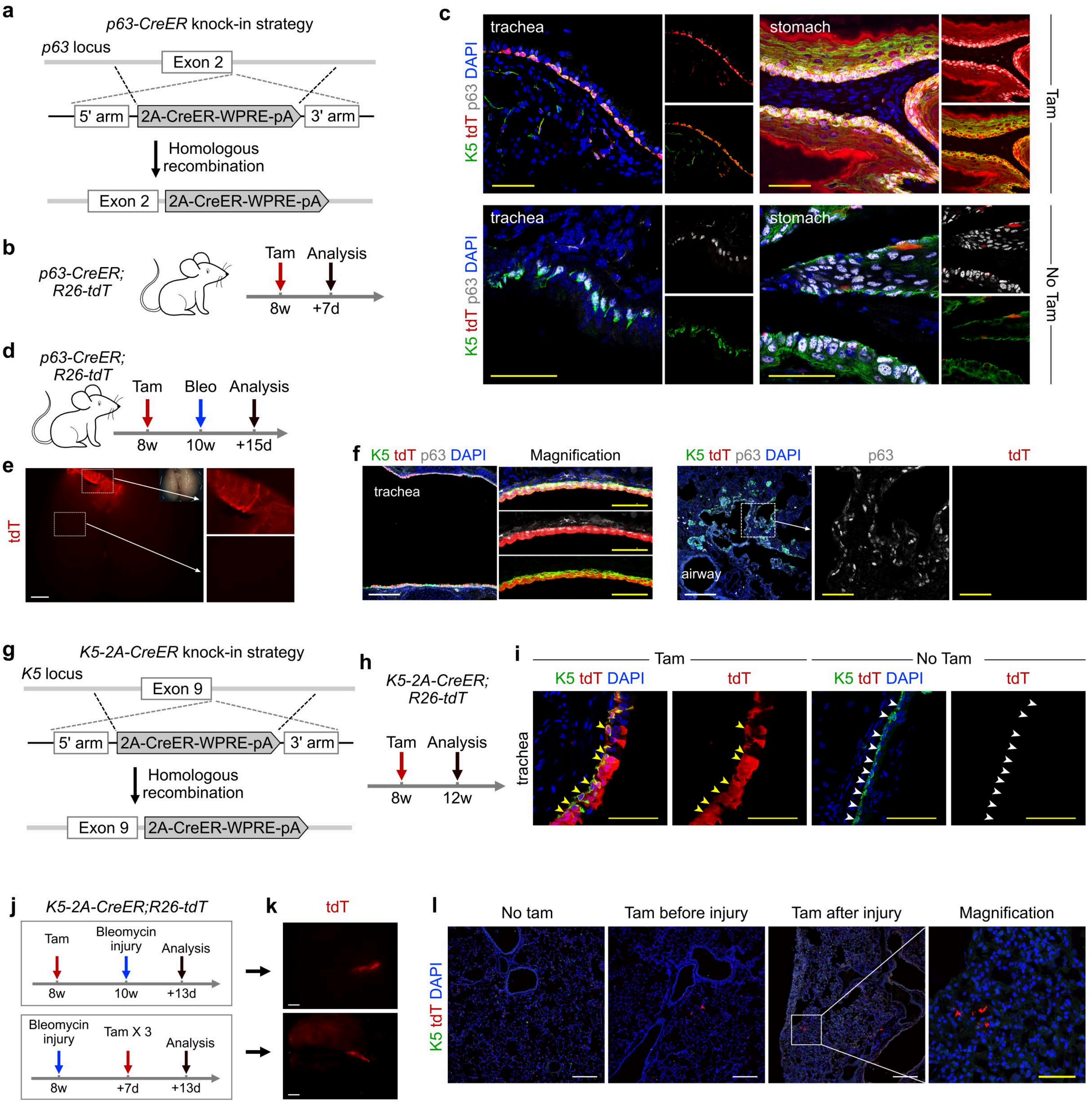


**Extended Data Fig. 1 | Lineage tracing of p63+ or K5+ cells in lung. a**, A schematic showing knock-in strategy of *p63-CreER* mice line. **b**, A schematic showing the experimental design. **c**, Immunostaining for K5, tdT, and p63 of tissue sections. Scale bar, 50 μm. **d**, A schematic showing the experiment design. **e**, Whole-mount fluorescent view of *p63-CreER;R26-tdT* lungs. Scale bar, 2mm. **f**, Immunostaining for K5, tdT, and p63 on tissue sections. **g**, A schematic showing knock-in strategy of *K5-2A-CreER* mice line. **h**, A schematic showing the experimental design. **i**, Immunostaining for K5, tdT, and p63 on tissue sections. Scale bar, 50 μm. **j**, A schematic showing the experiment design. **k**, Whole-mount fluorescent view of *K5-2A-CreER;R26-tdT* lungs. Scale bar, 2mm. **l**, Immunostaining for K5 and tdT on tissue sections. Scale bar, 200 μm. Each image is representative of 5 individual biological samples.

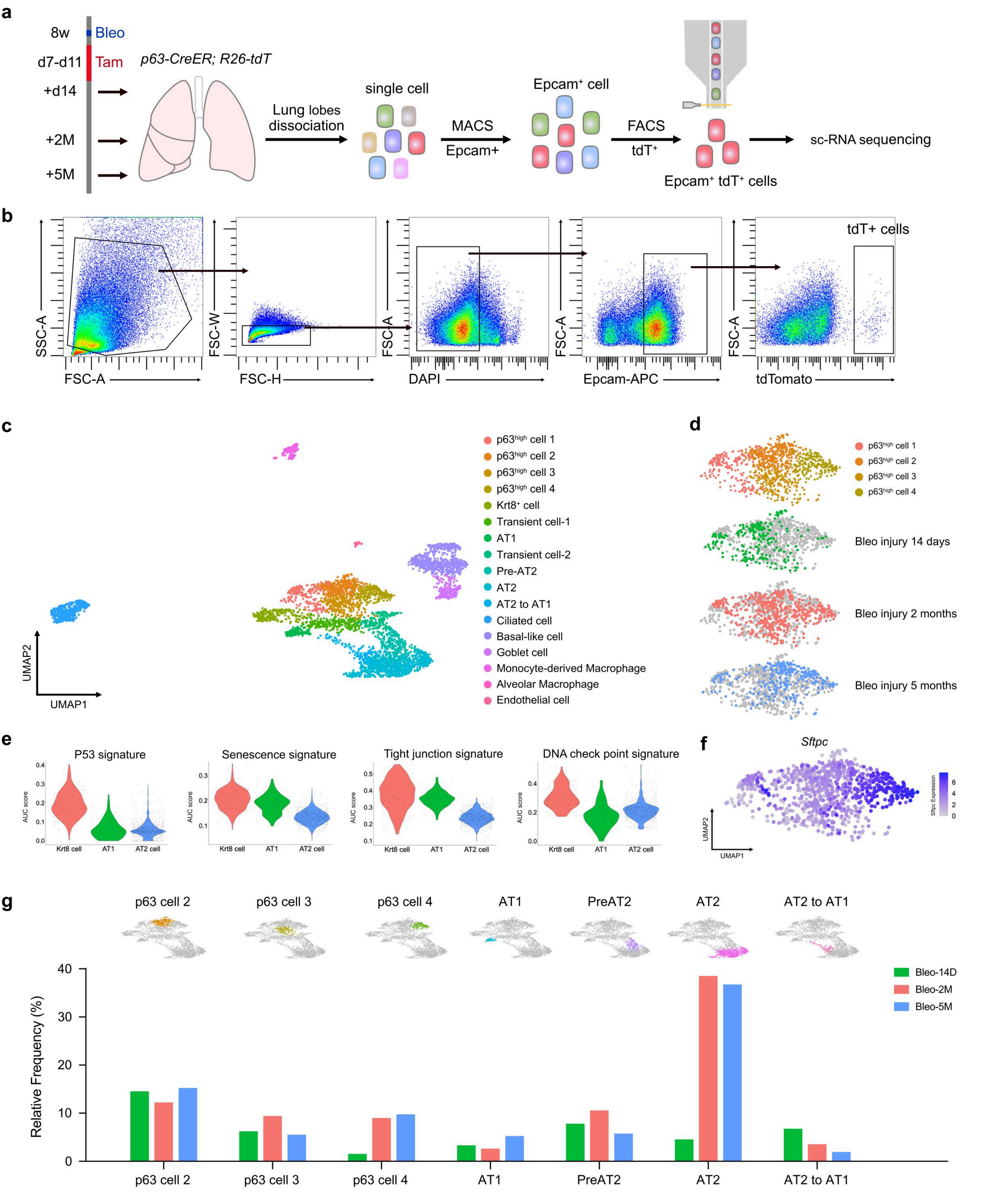


**Extended Data Fig. 2 | scRNA-seq analysis of p63⁺ progenitors. a**, A schematic showing the experiment design for isolating the p63⁺ cell lineage at 14 days, 2 months, and 5 months post injury. Epcam⁺ tdT⁺ live cells were performed by 10X sc-RNA sequencing process. **b**, Gating strategy of isolating p63⁺ cell lineages by FACS. **c**, UMAP visualization of each cell population isolated from bleomycin-treated lungs. **d**, UMAP plots showing the distribution of the p63$^{high}$ cells (top) and the samples collected at different stages (lower panels). **e**, Violin plots shows PATS-related signatures in the indicated cell populations from bleomycin-treated lungs. The black dots represent cells and the violin bodies indicate the distribution of these cells. **f**, UMAP plots showing *Sftpc* expression in p63$^{high}$ cells in bleomycin-treated lungs. **g**, UMAP plots showing the distribution of different cell types (top) and bar plots showing the relative frequency of the indicated cell types relative to all other cells at the indicated time points after injury (bottom).

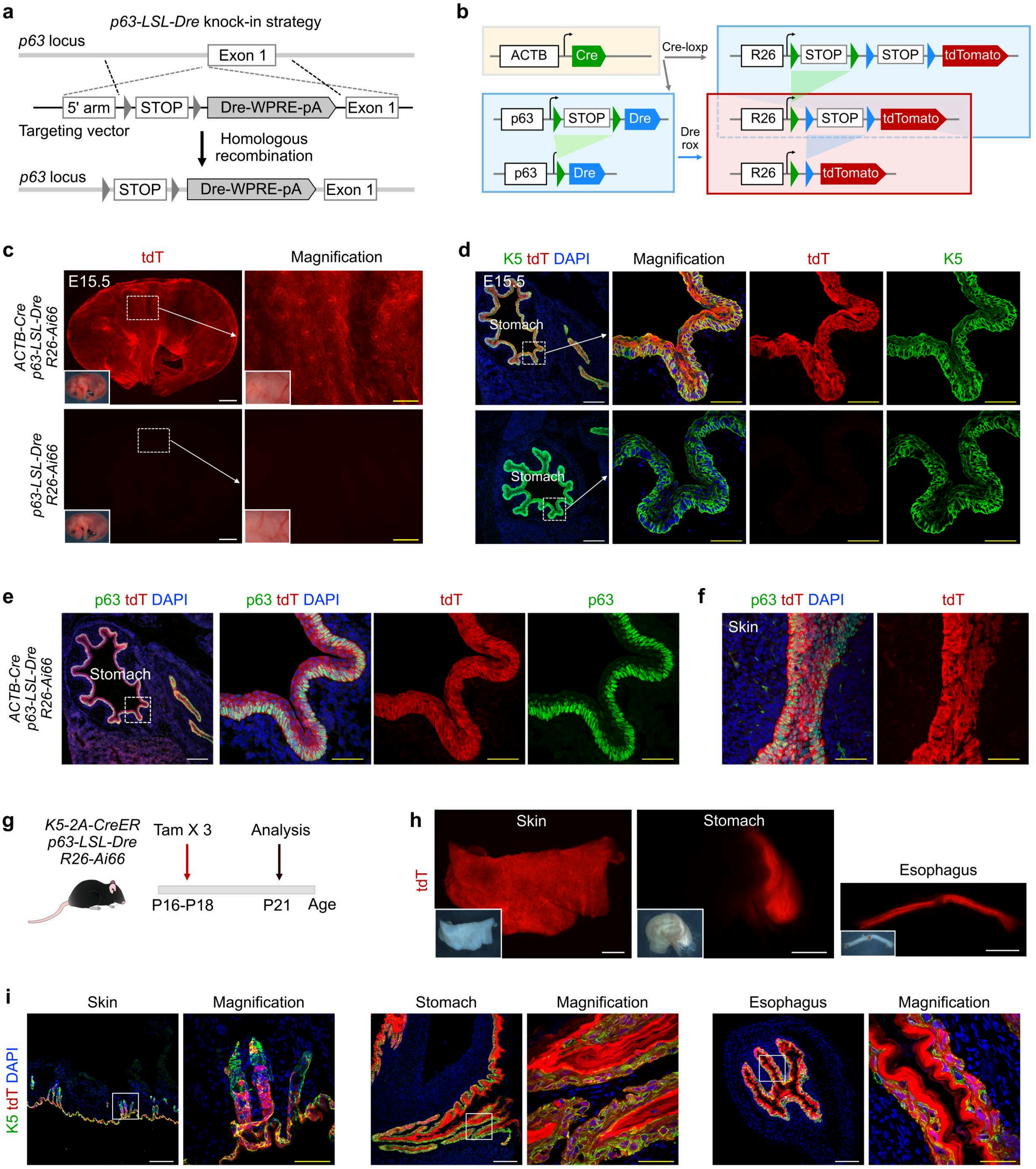


**Extended Data Fig. 3 | Generation and characterization of *p63-LSL-Dre* mouse. a**, A Schematic showing knock-in strategy for *p63-LSL-Dre* mouse generation. **b**, Mice crossing strategy for characterization of *P63-LSL-Dre* line. **c**, Whole-mount fluorescent view of *ACTB-Cre;p63-LSL-Dre;R26-Ai66* and *p63-LSL-Dre;R26-Ai66* embryos. Scale bars: white, 2 mm; yellow, 500 µm. **d**, Immunostaining for K5 and tdT on tissue sections. **e**,**f**, Immunostaining for K5, p63, and tdT on stomach and skin sections. **g**, A schematic showing the experimental design. **h**, Wholemount fluorescent view of organs collected from *K5-2A-CreER;p63-LSL-Dre;R26-Ai66* mice. Scale bars, 2 mm. **i**, Immunostaining for K5 and tdT on tissue sections. Scale bar in d,e,f,i: white, 200 µm, yellow, 50 µm.

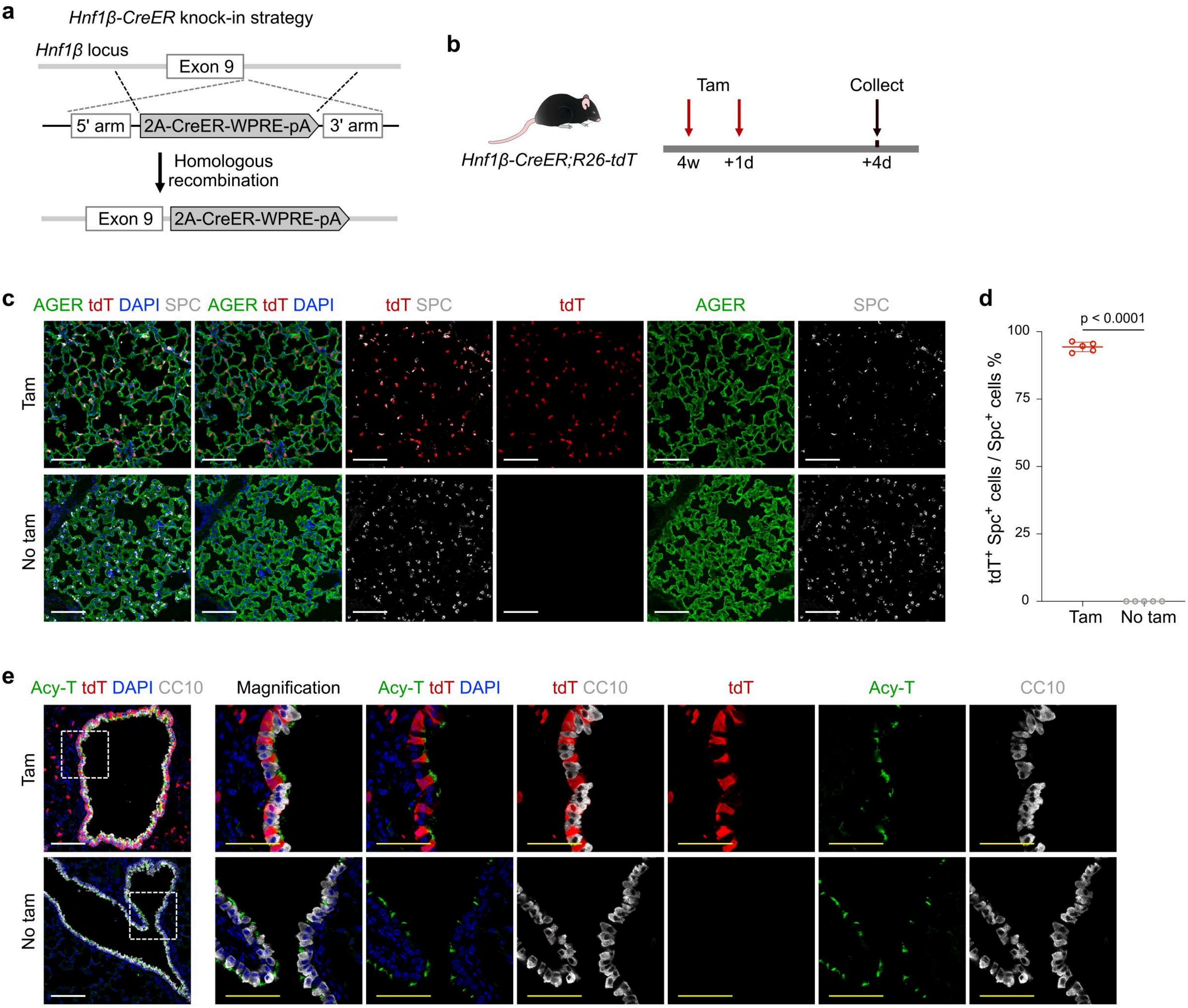


**Extended Data Fig. 4 | Generation and characterization of *Hnf1β-CreER* mouse.** **a**, A schematic showing the knock-in strategy for *Hnf1b-CreER* line. **b**, A schematic showing the experimental strategy. **c**, Immunostaining for AGER, tdT, and SPC on lung sections collected from *Hnf1b-CreER;R26-tdT* mice treated with or without Tam. **d**, Quantification the percentage of AT2 cells expressing tdT. n = 6 for each group. Two tailed student's t-test was used. **e**, Immunostaining for Acetylated-Tubulin, tdT, and CC10 on lung sections. Scale bar, 100 μm.

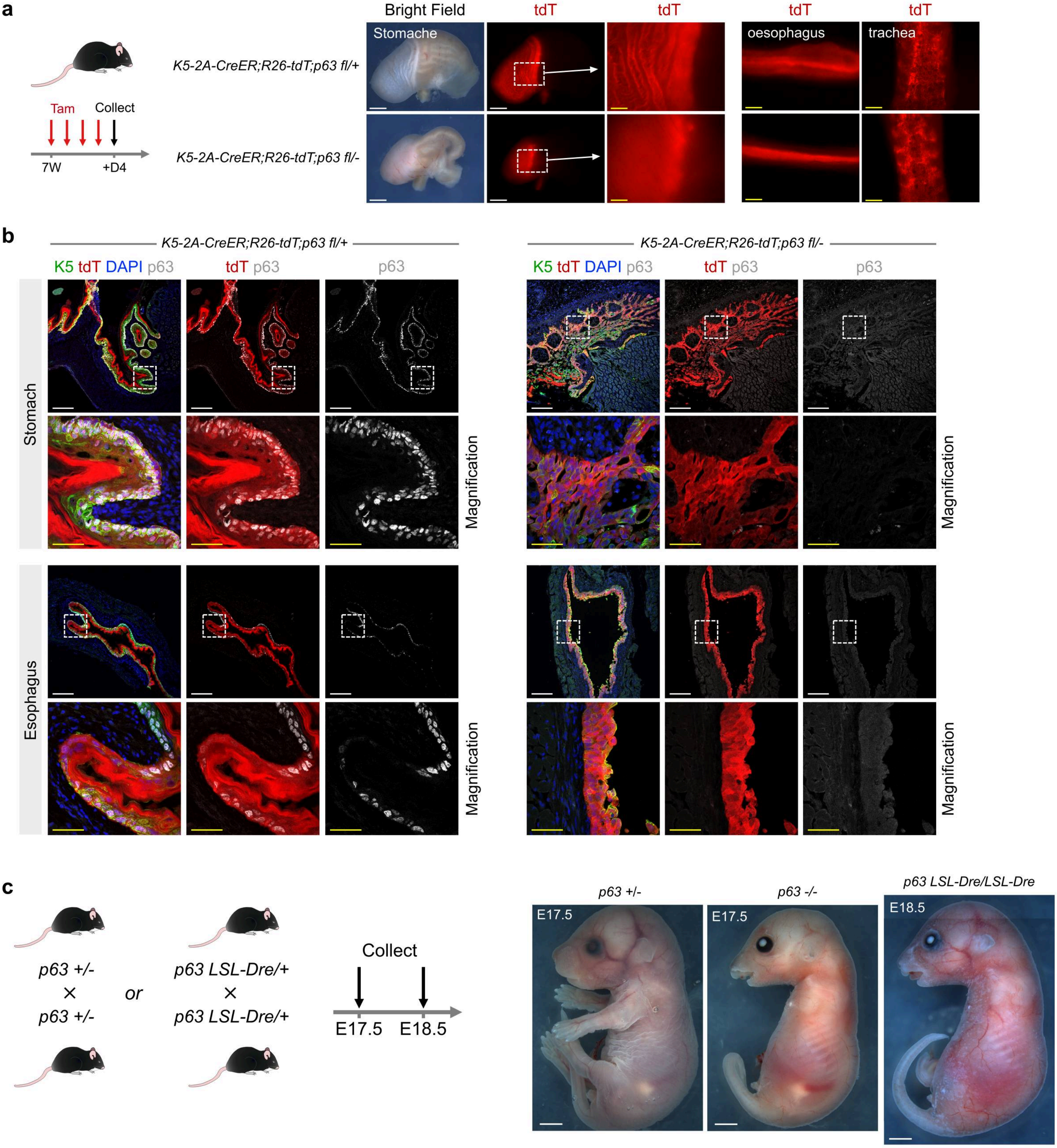


**Extended Data Fig. 5 | Characterization of *p63-fl* and *p63-LSL-Dre* mice.** **a**, Whole-mount fluorescence view of tissues collected from *K5-2A-CreER;R26-tdT;p63 fl/+* and *K5-2A-CreER;R26-tdT;p63 fl/–* mice. Scale bar, white, 2mm; yellow, 500 μm. **b**, Immunostaining for K5, tdT, and p63 on tissue sections. Scale bar, white, 200 μm; yellow, 50 μm. **c**, A schematic showing mice crossing strategy for generation of homozygous *p63 -/-* or *p63 LSL-Dre/LSL-Dre* mice (left panel). Right panel shows gross view of E17.5 or E18.5 p63 heterozygous or homozygous embryos. Scale bar, 2mm.

**Supplementary Table 1. Primer sequences for genotyping**

| Mouse line | Primer sequences | Product Size |
|---|---|---|
| *R26-tdTomato* | Forward: GGCATTAAAGCAGCGTATCC<br>Reverse: CTGTTCCTGTACGGCATGG | Mut: 196bp, WT: no band |
| | Forward: AAGGGAGCTGCAGTGGAGTA<br>Reverse: CCGAAAATCTGTGGGAAGTC | WT: 297bp, Mut: no band |
| *R26-Ai66* | Forward: ACGGGTGTTGGGTCGTTTGTTC<br>Reverse: ATGTTTCAGGTTCAGGGGGAGGTG | Mut: 404bp, WT: no band |
| | Forward: ACCATCGTGGAACAGTACGAGC<br>Reverse: TTGGTCACCTTCAGCTTGGC | Mut: 251bp, WT: no band |
| *ACTB-Cre* | Forward: CCTGGAAAATGCTTCTGTCCG<br>Reverse: CAGGGTGTTATAAGCAATCCC | Mut: 391bp, WT: no band |
| *p63-2A-CreER* | Forward: TCATGCCCTGAGCACATTGTAC<br>Reverse: TTCTTGCGAACCTCATCACTCG | Mut: 389bp, WT: no band |
| *p63-LSL-Dre* | Forward: TGCGCGGGACGTCCTTCTGCTAC<br>Reverse: CTCTACTTCCGCTGCTGCTCTTAT | Mut: 825bp, WT: no band |
| | Forward: GAGGCACCTGAATTCTGTTATCTT<br>Reverse: CTCTACTTCCGCTGCTGCTCTTAT | WT: 488bp, Mut: no band |
| *p63-DreER* | Forward: CTATTGCTTCCCGTATGGCTTTCA<br>Reverse: GTGTTGTAGGGGCTGGTGGACGAG | Mut: 806bp, WT: no band |
| | Forward: ACAGCCACAGTACACGAACC<br>Reverse: GCATATACAGCACCTCCTAA | WT: 518bp, Mut: no band |
| *Sox2-CreER* | Forward: AGGACTCGTGTTTGGGAACC<br>Reverse: CGCCGCATAACCAGTGAAAC | Mut: 1384bp, WT: no band |
| *CC10-CreER* | Forward: ACTCACTATTGGGGGTGTGG<br>Reverse: CCAAAAGACGGCAATATGGT | Mut: 245bp, WT: no band |
| | Forward: ACTCACTATTGGGGGTGTGG<br>Reverse: AGGCTCCTGGCTGGAATAGT | WT: 550bp, Mut: no band |
| *K5-2A-CreER* | Forward: CGGGCTCTACTTCATCGCAT<br>Reverse: ACCAAAGCATGTGGTTCTGC | Mut: 223bp, WT: no band |
| *Spc-CreER* | Forward: TGCTTCACAGGGTCGGTAG<br>Reverse: ACACCGGCCTTATTCCAAG | Mut: 210bp, WT: no band |
| | Forward: TGCTTCACAGGGTCGGTAG<br>Reverse: CATTACCTGGGGTAGGACCA | WT: 327bp, Mut: no band |
| *Hnf1β-2A-CreER* | Forward: CCTGGCCATTCCATCCAAGT<br>Reverse: GTTGCATCGACCGGTAATGC | Mut: 556bp, WT: no band |
| *p63-fl* | Forward: GCTTGAAGCATTGTTTCCTGT<br>Reverse: GTGGCACATGTCAACTTTGT | WT: 250bp, mut: 369bp |

Primer sequences for PCR identification of LSL recombination in *p63-LSL-Dre*

| Mouse line | Primer sequences | Product Size |
|---|---|---|
| *p63-LSL-Dre* | Forward: TCCATTGGAGTGGAGGAGCC<br>Reverse: CTGGTACTCCTTGCCGATGTT | full length: 2197bp, cut of LSL: 426bp |
